# THE DYNAMIC INTERPLAY BETWEEN HOMEODOMAIN TRANSCRIPTION FACTORS AND CHROMATIN ENVIRONMENT REGULATES PRONEURAL FACTOR OUTCOMES

**DOI:** 10.1101/2020.12.02.398677

**Authors:** Cheen Euong Ang, Victor Hipolito Olmos, Bo Zhou, Qian Yi Lee, Rahul Sinha, Aadit Narayanaswamy, Moritz Mall, Kirill Chesnov, Thomas Sudhof, Marius Wernig

**Affiliations:** Department of Bioengineering; Department of Pathology; Institute of Stem Cell and Regenerative Medicine; Department of Molecular and Cellular Physiology and Howard Hughes Medical Institute Stanford University, Stanford, CA 94305, USA

## Abstract

Generation of neurons of vast diversity involves early spatial and temporal patterning of the neuronal precursors by morphogenic gradients and combinatorial expression of transcription factors. While the proneuronal function of the basic-helix-loop-helix (bHLH) transcription factor Ngn2 is well established, its role in neuronal subtype specification remains unclear. Here, we found that coexpressing NGN2 with the forebrain homeobox factor EMX1 converts human pluripotent stem cells into a highly homogeneous glutamatergic forebrain neurons without partial cholinergic and monoaminergic gene programs observed in cells infected with NGN2 only. Our molecular characterization revealed that transcriptional output and genomic targeting of Ngn2 is altered by co-factors such as EMX1 explaining the more focused subtype specification. Ngn2 function is less modified by the chromatin environment and does not affect regionalization of pre-patterned neural progenitors. These results enable improved strategies for generating a plethora of defined neuronal subpopulations from pluripotent stem cells for therapeutic or disease-modeling purposes.

**Highlights:**

- NGN2 converts human ES cells into glutamatergic neurons some of which co-express a partial cholinergic program
- NGN2 directly binds to and activates ISL1 in ES cells which together with PHOX2A/B induce cholinergic genes
- Anterior-posterior regionalization affects NGN2 binding and transcriptional output but does not focus subtype specification
- Forebrain homeobox factors including EMX1 and FOXG1 redirect NGN2 chromatin binding and repress posterior and cholinergic genes, resulting in homogeneous forebrain excitatory neurons

## INTRODUCTION

The mammalian nervous system is the most diverse organ, comprising a plethora of neurons and glial cells organized along anterior-posterior and dorsal-ventral axes. Those neurons and glia cells differentiate from progenitor cells endowed each with a positional identity by spatially and temporally defined morphogenic gradients (Allan and Thor, 2015; Guillemot, 2007). It remains intriguing how mechanistically a handful of transcription factors and morphogens can give rise to the vast large neuronal diversity. One possibility is that the various proneural basic helix-loop-helix (bHLH) factors which are responsible to induce neuronal fates in progenitor cells assume different roles in subtype specifications (Aydin et al., 2019; Parras et al., 2002). For example, Ngn2 and Ascl1 often have mutually exclusive expression patterns, generating neurons of different subtypes such as glutamatergic and GABAergic neurons in the forebrain and spinal cord. In the retina, Ascl1/Math3 and NeuroD1/Math3 are important in the development of bipolar cells and amacrine cells, respectively (Inoue et al., 2002). Another possibility is that proneural factors do not possess subtype-specification potential and need to partner with cofactors, such as homeodomain and POU domain transcription factors, in different stages of neuronal differentiation (Guillemot, 2007). One of the well studied areas is the dorsal-ventral axis specification of the developing spinal cord. The progenitors along the dorsal-ventral axis in the spinal cord are marked by different homeodomain transcription factors, specifying the progenitors destined to give rise different neuronal subtypes. In postmitotic neurons, homeodomain transcription factors, such as Mnx1 and Crx, are crucial in motor neurons and cone cells differentiation (Freund et al., 1997; Jessell, 2000). Such transcription factors also function outside the context of normal development as combinations between proneural and lineage-specific factors can even convert fibroblasts to induced neuronal (iN) cells with different neuronal subtype specification (Ang and Wernig, 2014; Caiazzo et al., 2011; Pfisterer et al., 2011; Son et al., 2011; Tsunemoto et al., 2018; Yang et al., 2017).

We previously developed a protocol to rapidly and robustly generate functional neurons by NGN2 overexpression from human embryonic stem (hES) of induced pluripotent stem (iPS) cells. Those NGN2-iN cells express pan-neuronal and excitatory neuronal markers, fire repetitive action potentials and form functional synapses, making them a versatile platform to study cell biological processes in human neurons (Ang et al., 2019; Chanda et al., 2019; Konermann et al., 2018; Nehme et al., 2018; Pak et al., 2015; Yi et al., 2016; Zhang et al., 2013). It has been previously reported that NGN2 activity is context dependent: Overexpression of NGN2 in neural tube cell cultures under a low BMP condition promoted sensory fate while under a high BMP condition promoted an autonomic fate (Perez et al., 1999). It is thus unclear why overexpression of NGN2 which is widely expressed in the neural progenitors that give rise to neurons with different neurotransmitter subtypes (glutamatergic, cholinergic and noradrenergic neurons) only generates glutamatergic iN cells when expressed in hES/iPS cells. Here, we explored NGN2’s molecular chromatin function and its ability to induce specific neuronal programs in the context of different cell states and different transcription factor combinations using a series of single cell and bulk genomic sequencing techniques and epigenomic methods.

## RESULTS

### NGN2 induces three kinds of glutamatergic neurons in hES cells characterized by a partial cholinergic program

We have shown previously that forced expression of NGN2 in human ES and iPS cells results in an efficient conversion into functionally homogeneous excitatory neurons expressing markers of dorsal forebrain identity (Zhang et al., 2013). To better characterize the cell composition of these NGN2-iN cells, we performed single cell RNA-sequencing (scRNA-seq) using the smart-seq2 protocol at day 4 and 28 post NGN2 induction (**Figure 1A**). After filtering for cells with expression of at least 2,500 genes and with 200,000 paired end reads, we obtained 27 and 62 high quality cells for the day 4 and 28 timepoint, respectively (**Figure 1B**). We also obtained scRNA-seq data from 11 hES cells and included them into our analysis (Chu et al., 2016). When we performed principal component analysis (PCA), the cells separated into three clusters largely corresponding to cells from the three time points with the first principal component corresponding to genes enriched for gene ontology (GO) terms associated with cell proliferation (*DNA replication, cell division and G1/S transition of cell cycle*) and the second principal component corresponding to genes enriched in nervous system development (**Figure 1C, Figure S1A**). This reflects our previous observation that NGN2 drives hES rapidly out of cell cycle and then neuronal differentiation and maturation occurs over several weeks (Chanda et al., 2019; Zhang et al., 2013). When we performed hierarchical clustering of genes that are four-fold changed among the different time points, we found that ES specific genes (*POU5F1, NANOG, SOX2*) were downregulated precipitously in the d4 and d28 timepoints. As expected, transcription factors reported to be downstream of NGN2 (*NEUROD1, NHLH1, NEUROD4 and HES6*) were upregulated at day 4 and subsequently downregulated. Mature neuronal markers (*MYT1L*) and neuronal subtype markers (*ISL1, PHOX2B*) were induced and maintained during differentiation (**Figure 1D, Figure S1B**). To assess the degree of heterogeneity of day 4 and 28 iN cells, we performed t-distributed stochastic neighbor embedding (tSNE) using genes that are most variable across hES, day4 iN and day28 iN cells (y_cutoff_=0.75, x_cutoff_=0.5, variable genes=1144). We found that day 28 iN cells homogeneously express glutamatergic markers [*VGLUT2* (S*LC17A6), VGLUT1 (SLC17A7*)] confirming our previous findings that the NGN2 hES-iN cells contain exclusively glutamatergic functional synapses (Zhang et al., 2013). However, the day 28 iN cells formed three relatively distinct clusters with similar cell numbers (**Figure 1E, F, G**). One cluster of cells was positive for cholinergic transporters [*vAChT* (*SLC18A3), ChT (SLC5A7)*] and the two transcription factors, *ISL1 and PHOX2B*. Another cluster was characterized by glutamatergic genes and *ISL1*, but not *PHOX2B* expression. The third cluster expressed glutamatergic markers only (**Figure 1F, H**). To validate the results of the scRNA-seq data, we performed immunofluorescence for ISL1 and PHOX2B at day 4. Immunofluorescence data confirmed the presence of ISL1+ cells among the FLAG-NGN2 infected cells (~50%, n=3). 25% of the ISL1 positive cells were also PHOX2B positive (**Figure 1I, J, K, L**). These data demonstrate that NGN2 induces exclusively glutamatergic neurons, but a subset co-expresses cholinergic markers. Importantly, the induction of a cholinergic subtype is only partial, since CHAT, the rate limiting enzyme for acetylcholine synthesis, is not expressed in most cells (**Figure S1D**).

**Fig. 1.**
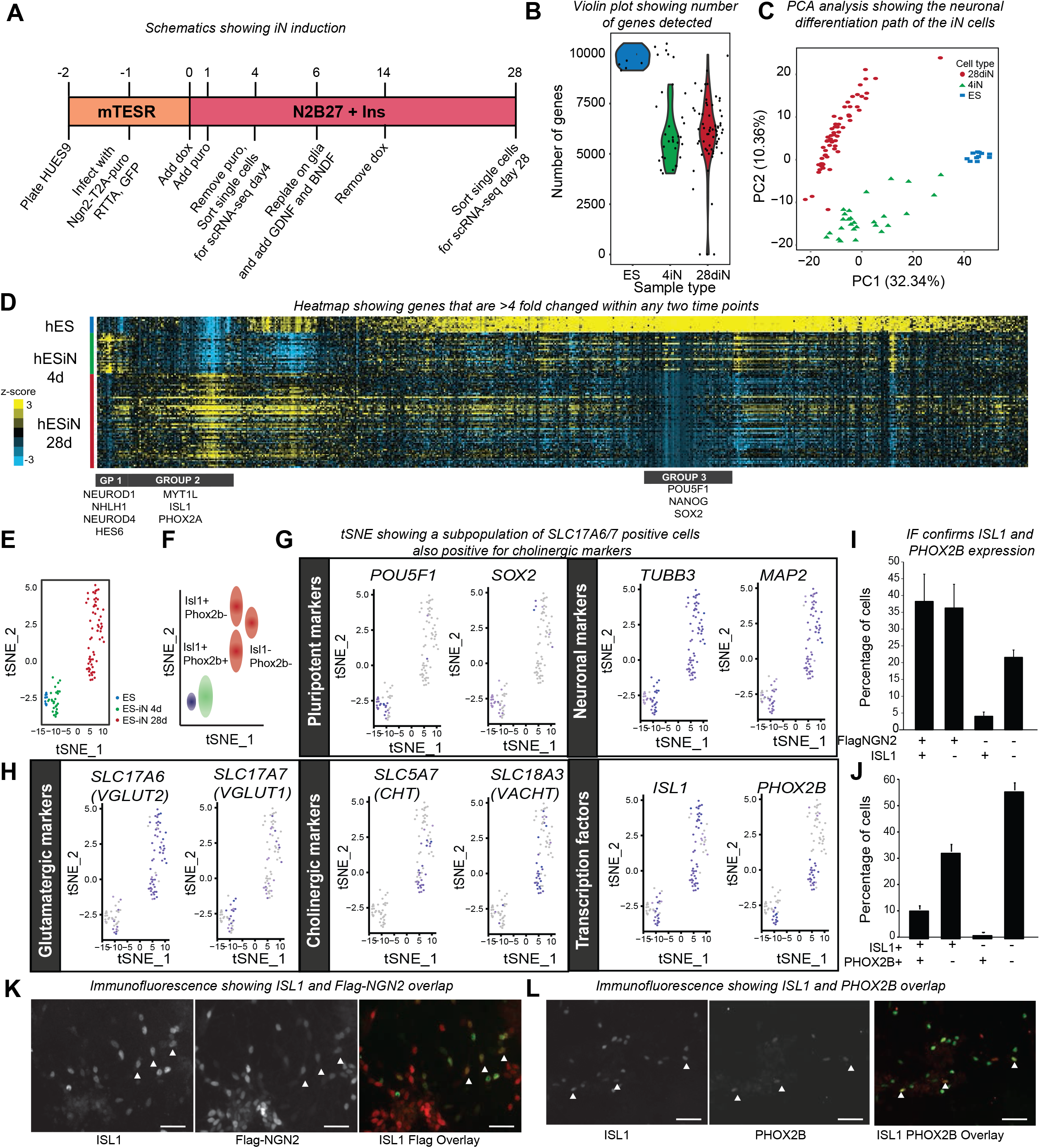
Single cell RNA-sequencing reveals heterogeneity of Ngn2 iN cells. (A) Differentiation protocol to obtain the day 4 and 28d NGN2 induced neuronal (iN) cells. Day 4 represent immature postmitotic neurons and day 28 represent mature neurons. In total, we obtained 27 day 4 and 62 day 28 high quality iN cells. (B) Violin plot showing the number of genes detected by single cell RNA-sequencing in undifferentiated ES cells, day 4, and day 28 iN cells. Only cells that had 200,000 reads and more than 2,500 genes were considered for subsequent analysis. The ES cell scRNA-sequencing (11 cells) was downloaded from NCBI GEO (GSM1964970). (C) Principal component analysis showing the progression of iN cell reprogramming. The gene ontology analysis of genes corresponding to the first and second principal component are shown in Figure S1A. (D) Hierarchical clustering of genes that are changed at least four-fold among the three time points. Group 1 contain direct NGN2 target genes which tend to be highest in day 4 cells. Group 2 includes mature neuronal markers. Group 3 genes include pluripotency markers. The GO terms for each cluster are listed in Figure S1B. (E) T-distributed stochastic neighbor embedding (t-SNE) plot for the three different cell populations: ES cells (blue), NGN2 4d (green) and 28d (red) iN cells plotted using Seurat with the following settings (y_cutoff_=0.75, x_cutoff_=0.5, variable genes=1144) (F) Key for specific cell populations described in Figure 1G and 1H. (G) t-SNE plot for two ES cell markers (POU5F1 and SOX2) and neuronal markers (TUBB3 and MAP2). Grey and purple representing low and high expressing cells respectively. (H) t-SNE plot for two markers for glutamatergic (SLC17A6 and SLC17A7) and cholinergic neuronal markers (SLC5A7 and SLC18A3). The day 28 cells that are positive for both cholinergic markers are also positive for ISL1 and PHOX2B. Grey and purple representing low and high expressing cells respectively. (I) Quantification of day 28 NGN2 iN cells for FLAG (Flag-NGN2) and ISL1 immunofluorescence. Double- and single-positive, and double-negative cells are shown as percentage of all DAPI positive cells (n=3). Error bars = s.e.m. (J) Quantification of ISL1 and PHOX2B positive cells as in I (n=3). Error bars = s.e.m. (K) Immunofluorescence images of ISL1 (Left) and FLAG (Middle) and the overlay (Right). White triangles: ISL1:FLAG double positive cells. Scale bar: 50μm (L) Immunofluorescence images of ISL1 (Left) and PHOX2B (Middle) and the overlay (Right). White triangles: ISL1 and PHOX2B double positive cells. Scale bar: 50μm

ISL1 and PHOX2B are known to be expressed in spinal/cranial motor neurons and sympathetic/parasympathetic neurons (Ericson et al., 1992; Pattyn et al., 1997). Notably, their overexpression induces cranial or spinal cholinergic neurons (Mazzoni et al., 2013). Moreover, *ISL1, PHOX2B, SLC18A3 and ChT (SLC5A7)* are not detected by scRNA-sequencing in neurons and glial cells in human cortex (medial temporal gyrus) (**Figure S1G**). We therefore found the prominent ISL1 induction following NGN2 expression surprising and asked whether the level of NGN2 transgene expression may affect ISL1 expression in the hES cell system. However, we found no correlation between the intensity of FLAG staining (indicative of NGN2 level) to that of ISL1 (R^2^ = 0.0002, Pearson = −0.01) indicating that NGN2 may induce ISL1 only indirectly (**Figure S1E**). Similarly, little to no correlation was found between ISL1 and PHOX2B expression levels (R^2^ = 0.0534, Pearson = 0.23) but there is a Boolean relationship with PHOX2B+ cells being a subset of the ISL1+ cell population (**Figure 1L, S1F**).

A closer examination on the neurotransmitter subtype specific transporters or rate limiting enzymes for GABAergic, monoaminergic and cholinergic neurons showed that the transporters and enzymes required for the neurotransmitter production and release are not all expressed within the same cell (**Figure S1C, D**) signifying that the induction of subtype specification genes is not complete. This intriguing finding raises the possibility that NGN2 induces multiple neurotransmitter subtype specific gene programs and additional mechanisms must complement NGN2 to accomplish precise subtype-specification (**Figure S1D**).

### Cholinergic gene induction in hES cells by NGN2 is mediated by ISL1 and PHOX2B

Our scRNA sequencing data showed a correlation between ISL1 and PHOX2B and the cholinergic gene program. We wondered whether ISL1 and PHOX2B are responsible for the induction of cholinergic genes initiated by NGN2 in hES cells (**Figure 2A**). To that end, we overexpressed ISL1, PHOX2B and both genes together with NGN2 and found that all transcription factor combinations produced β-III-tubulin positive cells as early as day 4 (**Figure 2B**). Quantitative RT-PCR on day 4 iN cells showed that ISL1 and PHOX2B induced expression of two cholinergic genes (*vAChT (SLC18A3)* and *ChT (SLC5A7)*) (**Figure 2D**). CHAT was only induced moderately and only by ISL1 (**Figure 2D**). The same findings were reproducible in another ES cell line (**Figure S2C**). We confirmed induction of ChT (SLC5A7) on the protein level by Western blotting which also revealed that neither did ISL1 induce additional PHOX2B compared to control, nor did PHOX2B induce ISL1 suggesting that the induction of cholinergic genes by those two transcription factors is independent of each other (**Figure 2C**). Next, we asked whether ISL1 and PHOX2B would be necessary for induction of cholinergic genes. Using two hairpins specific to ISL1, we found that ISL1 downregulation indeed reduced the expression of CHAT, vAChT (SLC18A3) and ChT (SLC5A7) (**Figure 2E, G-H**). The downregulation of ChT (SLC5A7) could be confirmed by Western blotting (**Figure 2F**). Unlike ISL1, PHOX2B knock-down only reduced the induction of ChT (SLC5A7), the gene most prominently induced by PHOX2B (**Figure S2A, B**) while the other two cholinergic genes remained unchanged. In summary, in hES cells, NGN2 indirectly induces the expression of cholinergic genes via the initial induction of ISL1 and PHOX2B both of which induce cholinergic expression independently of each other. Of note, the ISL1 and PHOX2B induction occurs only in a subset of NGN2 infected cells and the cholinergic programs are incomplete (i.e. cells do not express all three cholinergic genes CHAT, ChT (SLC5A7), vAChT (SLC18A3) required for proper acetylcholine synthesis and release).

**Fig. 2.**
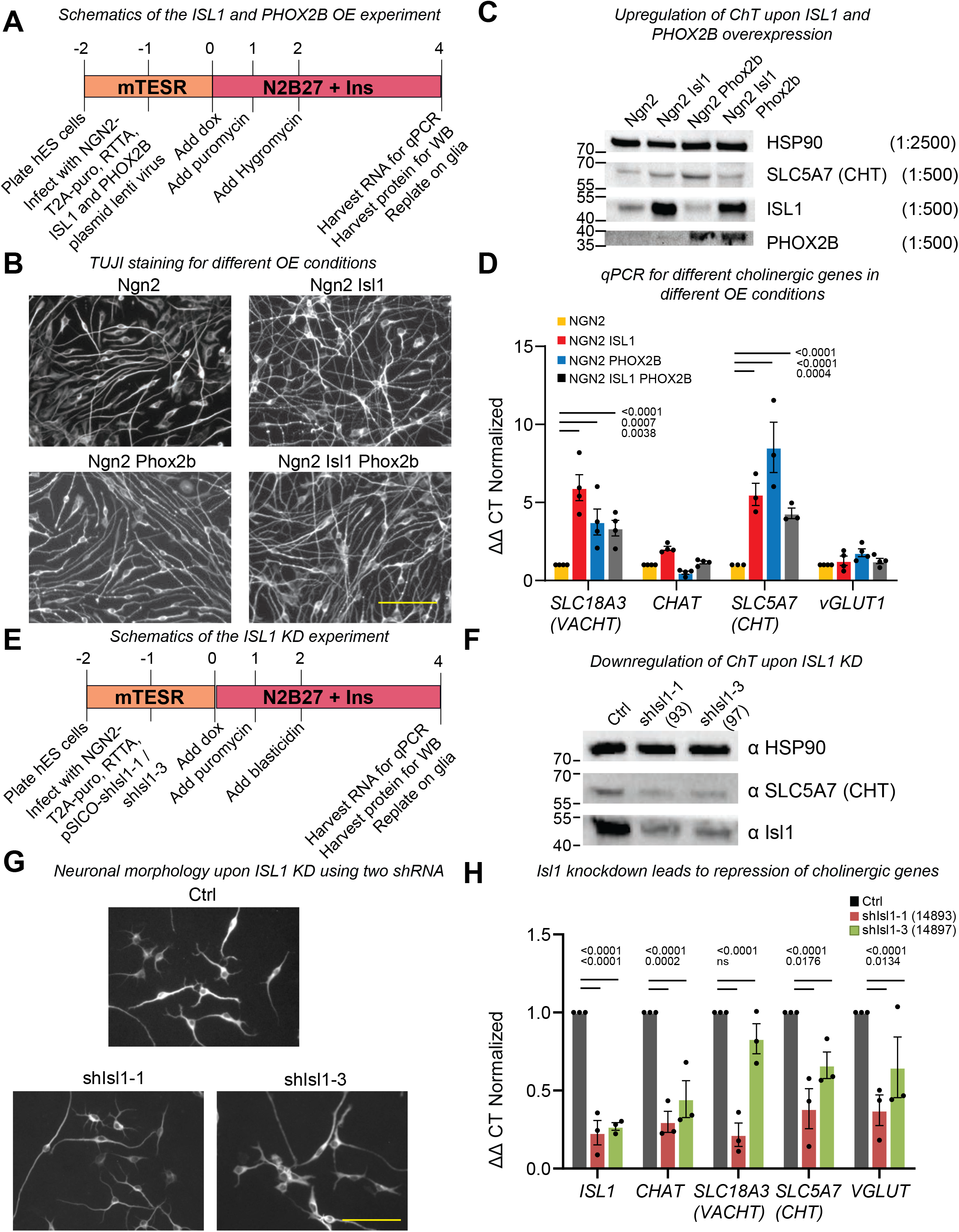
ISL1 and PHOX2B is sufficient to induce cholinergic genes. (A) Outline to generate day 4 Ngn2 iN cells with additional expression of ISL1, PHOX2B or ISL1:PHOX2B. (B) Immunofluorescence images of β-III-tubulin (TUJI) staining of day 4 Ngn2 iN cells alone or coexpressing ISL1, PHOX2B and ISL1 PHOX2B. Scale bar: 100μm (C) Western blot analysis of cells in day 4 NGN2 iN cells alone or coexpressing ISL1, PHOX2B, and ISL1:PHOX2B using antibodies indicated. (D) Quantitative RT-PCR examining the day 4 NGN2 iN cells alone or coexpressing ISL1, PHOX2B and ISL1:PHOX2B for three cholinergic markers (CHAT, SLC18A3, SLC5A7) and a glutamatergic marker (vGLUT1). (N=3, error bars = s.e.m. ANOVA. Exact adjusted p-values are marked in the graph) (E) Outline to generate day 4 iN cells expressing a control or two different ISL1 short hairpins. (F) Western blot analysis of day 4 iN cells expressing a control or two ISL1 hairpins probing for SLC5A7 (ChT), Isl1, and HSP90 as loading control. 14893 (93) and 14897 (97) denotes two different hairpins. (G) Morphology of day 4 Ngn2 iN cells infected with a control or two ISL1 hairpins. Shown is GFP immunofluorescence of GFP-expressing iN cells. Scale bar: 100μm (H) Repression of cholinergic genes by ISL1 knock-down as shown by qRT-PCR of day 4 Ngn2 iN cells infected with control or two ISL1 shRNAs. (N=3, error bars represent s.e.m. ANOVA. Exact adjusted p-values are shown in the graph). 14893 and 14897 denotes two different hairpins.

### Regionalization of donor cells is maintained throughout NGN2-mediated differentiation but does not resolve neuronal subtype blurring

During development, NGN2 is induced in neural progenitor cells after they have been endowed with a positional identity. Those neural progenitors with different positional identities subsequently differentiate into forebrain glutamatergic neurons, spinal cholinergic motor neurons and peripheral sensory neurons. This may suggest that NGN2 induces different subtype differentiation programs based on the developmental history and the positional identity of the neural progenitor cells (Brunet and Ghysen, 1999). Given that our hES-iN protocol bypasses the regional identity specification step, we hypothesized that NGN2 overexpression in neural progenitor cells with specific regional identities can help restrict the neurotransmitter subtype of the resulting iN cells such as the partial induction of a cholinergic program by limiting the promiscuous action of NGN2. To test this idea, we first differentiated hES cells into anterior (SL, treated with **S**B431542 and **L**DN193189) and posterior (SLC, treated with **S**B431542, **L**DN193189 and **C**HIR99021) neuroectodermal cells and then expressed NGN2 to induce their differentiation into neurons (**Figure 3A, Figure S3A**) (Chambers et al., 2009; Du et al., 2015). As a control we also included hES cells expressing NGN2 without prior neuralization and regionalization. We confirmed that anterior progenitors (SL) were positive for OTX2 and posterior neural progenitors (SLC) were positive for HOXA3 by immunofluorescence (**Figure 3B**) and qRT-PCR (**Figure S3C**). We picked anterior and posterior neuroectodermal cells since NGN2 generates glutamatergic cortical projection neurons in dorsal forebrain progenitors and induces cholinergic motor neurons in ventral progenitors of the developing spinal cord during mouse CNS development (Parras et al., 2002). We hypothesized that those drastically different starting populations would present NGN2 to different chromatin environments affecting neurotransmitter subtype specification.

**Fig. 3.**
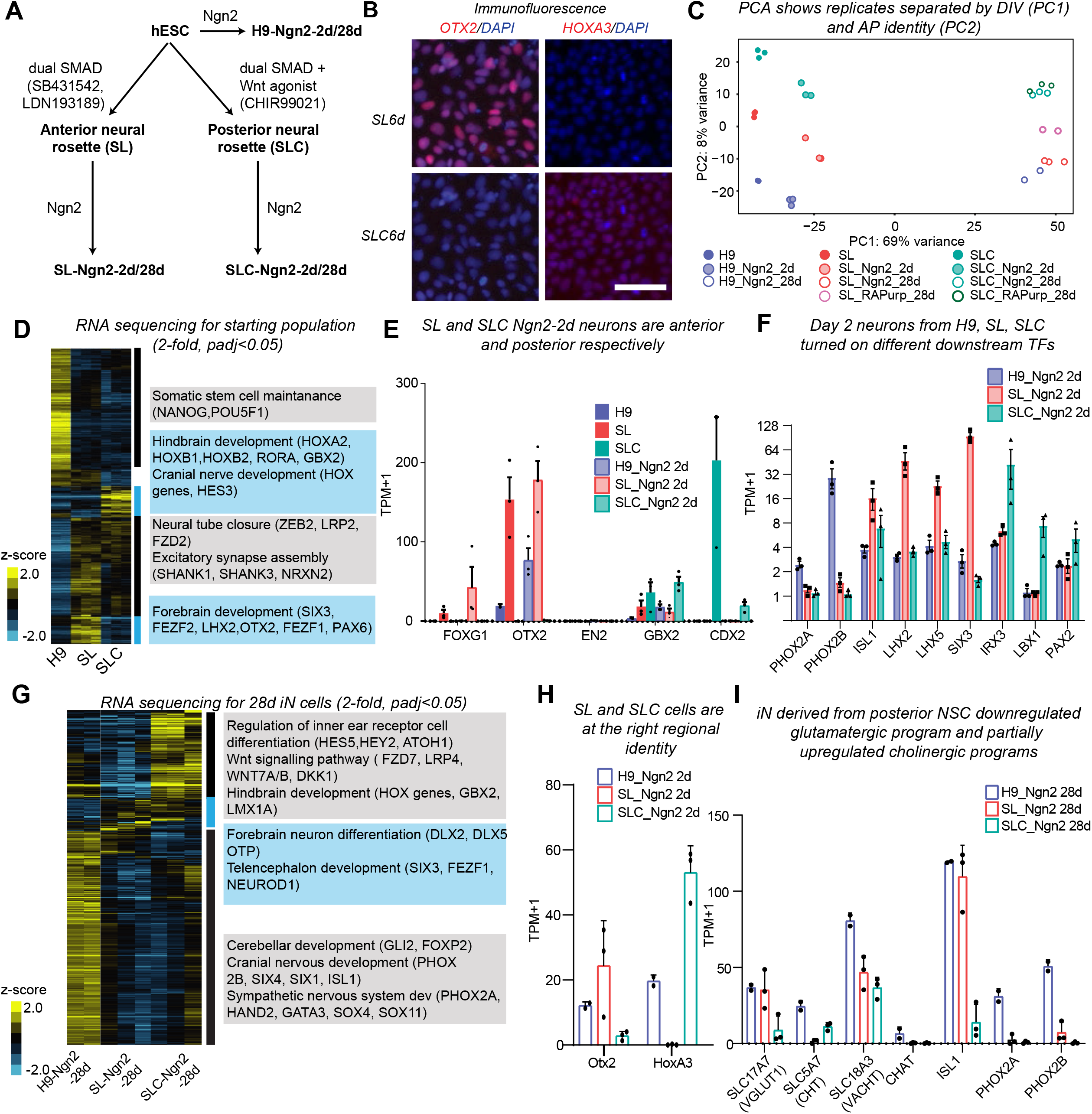
Regionalization of donor cells is maintained throughout Ngn2-induced neuronal differentiation. (A) Patterning of human ES cells into anterior neural cells with SB431542 and LDN193189 (SL) and posterior neural cells with SB431542, LDN193189, and the Wnt antagonist CHIR99021 (SLC). See also Figure S3A for details. (B) Immunofluorescence of OTX2 (an anterior marker) and HOXA3 (a posterior marker) validating the positional identity of most anterior and posterior neural cells at day 6 post differentiation. Scale bar: 50μm (C) Principal component analysis of RNA sequencing of the three donor cell populations: ES cells (H9), anterior (SL), and posterior (SLC) neural progenitors and their corresponding day 2 and 28 iN cells. The first principal component separates the samples by the differentiation stage and the second principal component separates the cells derived from different starting populations. (D) Hierarchical clustering of the genes (>=2-fold change and p adj<0.05) of the three starting populations [H9 (n=2), SL (n=3) and SLC (n=3)]. Significant gene ontology terms and the genes contributing to them for each highlighted cluster are listed in order of the highlighted area (PANTHER, p adj<0.05). (E) Expression of positional identity genes (FOXG1, OTX2, EN2, GBX2, CDX2) in the starting populations (H9, SL, SLC) and their corresponding day 2 iN cells. Error bars = s.e.m. (F) Expression of different downstream transcription factors in different starting populations. Error bars = s.e.m. (G) Hierarchical clustering of significant genes (>=2-fold change and p adj<0.05) of day 28 Ngn2 iN cells from the three starting populations [H9-NGN2-28d (n=2), SL-NGN2-28d (n=3), and SLC-NGN2-28d (n=3)]. Significant gene ontology terms and the genes contributing to them for each highlighted cluster are listed in order of the highlighted area (PANTHER, p adj<0.05). (H) Expression of positional identity genes (OTX2, HOXA3) in day 28 Ngn2 iN cells derived from different starting populations (H9, SL, SLC). Error bars = s.e.m. (I) Expression of a glutamatergic gene (SLC17A7), cholinergic genes (SLC5A7, SLC18A3, CHAT) and cholinergic transcription factors (ISL1, PHOX2A, PHOX2B) in day 28 Ngn2 iN cells derived from the three starting populations (H9, SL, SLC). Error bars = s.e.m.

All three starting populations [H9-hES, anterior (SL) or posterior (SLC) neural progenitor cells] produced mature iN cells with elaborate processes upon NGN2 induction (**Figure S3B**). To investigate how these different starting populations affect subsequent subtype specification, we performed RNA sequencing on the three starting populations (H9-hES, SL and SLC) and their corresponding day 2 iN cells (immature) and day 28 (mature) iN cells. Principal component analysis revealed an intriguing phenomenon: The stronger first principal component (explaining 69% of the variance) expectedly indicated the cell maturation (**Figure 3C**). However, the second, weaker principal component (explaining 8% of the variance) accounted for the different starting populations irrespective of differentiation stage (**Figure 3C**). This indicates that certain features of the starting cell population (and the most prominent different features between SL and SLC neuroectoderm is their regionalization) is not affected by NGN2 and remains throughout NGN2-mediated neuronal differentiation.

To further investigate this possibility, we inspected expression of region-specific genes. Analysis of differentially expressed genes among the three starting cell populations (padj<0.05, 2-fold) showed genes involved in forebrain development are enriched in the anterior population and genes for hindbrain and cranial development are enriched in the posterior population (**Figure 3D**). Classic anterior (*FOXG1 and OTX2*) and posterior (*CDX2*) marker genes were among the differentially expressed genes as expected with minimal expression of midbrain markers such as *EN2 and GBX2* (**Figure 3E**). To understand whether the regional identity of the pre-patterned neural progenitor cells is maintained during NGN2-mediated differentiation, we inspected differentially expressed genes among the three populations two days after NGN2 expression (padj<0.05, 2-fold). We found that the pre-established regional identities were preserved in these immature neuronal cells reflected in the rhombomere development GO term enrichment in the posterior cells and a forebrain regionalization GO term in the anterior cells (**Figure S3D**). This suggests that the NGN2-induction of neuronal specification does not overwrite the pre-established regional identities and that NGN2-iN cells can be endowed with more refined regional identities by patterning neural progenitor cells (**Figure S3D, 3H**). Nevertheless, we found that NGN2 induces different downstream transcription factors depending on region-specific chromatin configuration. In case of the day 2 NGN2 iN cells derived from anterior neural stem cells, anterior progenitor markers LHX2/5 and SIX3 are higher compared to that from H9 and the posterior neural stem cells; meanwhile for day 2 NGN2 iN cells derived from posterior neural stem cells, different spinal cord progenitor domain markers IRX3, LBX1 and PAX2 are higher compared to that from H9 and the anterior neural stem cells (**Figure 3F**). With respect to day 28 iN cells, GO term analysis and inspection of key region-specific genes of the RNA sequencing showed that the regional identity of day 28 iN cells was well maintained (**Figure 3G, H**). For example, telencephalic development genes (*SIX3, FEZF1*) were induced in SL-NGN2-28d iN cells and inner ear receptor cell genes (*HEY2, ATOH1*) were induced in day 28 SLC-NGN2-28d iN cells. While the H9-derived iN cells expressed genes involved in cerebellar, cranial and sympathetic neuron development, the iN cells derived from the two pre-patterned neural stem cells downregulated those genes precipitously (**Figure 3G**).

Next, we addressed our initial hypothesis and investigated the neurotransmitter subtype specification of day 28 NGN2-iN cells derived from the three different starting populations. Contrary to our expectation, the patterning did not fundamentally affect the induction of a cholinergic program which remained partial in all three conditions (**Figure 3I**). In particular the posterior condition yielded rather a decreased than increased expression of cholinergic genes and failed to induce CHAT. Western blot analysis confirmed noticeable decrease of CHT in SL-NGN2-28d (SLC5A7) protein level compared to the other two populations (**Figure S3F**). Although none of the three different conditions eliminated the cholinergic program completely or induced a full cholinergic subtype, quantitative expression differences were observed. For example, *ISL1* and *vAChT (SLC18A3)* is lowest in SLC-NGN2-28d, and *PHOX2A/B* as well as *ChT (SLC5A7)* are low in both SL-NGN-28d and SLC-NGN-28d compatible with our previous finding that *ChT (SLC5A7)* is most induced by PHOX2B and that vAChT *(SLC18A3)* is most induced by ISL1 (**Figure 2D, Figure 3I**).

Similar to cholinergic subtype specification, we found no clear pattern between the three groups when we plotted the expression of marker genes for different neurotransmitter phenotypes (**Figure S3E**). On the other hand, excitatory genes are prominently expressed in iN cells from all three cell types, although somewhat reduced in SL and SLC conditions (**Figure S3E**). Electrophysiology confirmed a glutamatergic specification since both SL-NGN2-28d and SLC-NGN2-28d iN cells exhibited exclusively miniature excitatory postsynaptic currents (EPSCs) which could be blocked by the AMPA receptor blocker CNQX (**Figure S3G**). The excitatory phenotype was further corroborated by homogeneous expression of the glutamatergic marker vGLUT1 as determined by immunofluorescence (**Figure S3H**).

To examine whether regionalizing stimuli after NGN2 induction would alter neurotransmitter subtype specification, we systematically added agonists and antagonists targeting major signalling pathways (TGFβ, BMP, Wnt, FGF, RA, SHH) after doxycycline induction for 7 days and examined the level of cholinergic genes of the H9-iN cells at day 7. We observed no significant upregulation or downregulation of ISL1, ChT (SLC5A7), VGLUT1 with the exception of CHIR99021 (Wnt agonist) and Trametinib (FGF antagonist) which induced immature/stressed neuronal cells and cell death, respectively (**Figure S3I**). Thus, regionalizing factors after neuronal induction has even less effect on neuronal subtype-specification than regionalizing the progenitor cells.

In summary, our findings showed that the regional identity of the starting cell population is stably inherited by neurons after NGN2 induction, and thus is unaffected by NGN2. At our level of sophistication, the broadly defined anterior and posterior differentiated progenitor populations used are insufficient to induce complete neurotransmitter subtype specification.

### Genomic binding of NGN2 is dependent on the chromatin configuration

We next sought to molecularly explain the context-specific effects of NGN2 and asked whether the chromatin state affects the physical binding of NGN2. The genomic binding of Ascl1, the other prominent proneural bHLH factor, was shown to be similar between cell types (Wapinski et al., 2013). We performed ChIP-sequencing of NGN2 and found 2625, 706 and 770 high-confidence peaks in H9, SL and SLC cells, respectively, two days after NGN2 induction. Removing the peaks that are also present in rtTA control, we obtained 2018 sites among H9-NGN2-2d, SL-NGN2-2d and SLC-NGN2-2d. Unexpectedly, while there was a large degree of overlap, NGN2 binding was most widespread when induced in undifferentiated ES cells and more restricted in differentiated population (SL and SLC) and clearly distinct between the three cell types (**Figure 4A**). The bHLH motif (CANNTG) was most significantly enriched in all cell types (E-values: 1.0E-676, 1.9E-682, 8.9E-319, respectively) (**Figure 4B, C**) with most peaks harboring more than one bHLH motif (**Figure S4A**). Analyzing different types of E-boxes we found a preference for CA**GA**TG E-box motif in H9 cells and a preference for CA**GC**TG motifs in the SL and SLC cells (**Figure S4B**). When we compared the peak classification in all three conditions, the majority of the peaks were located in intergenic and intronic regions and only a small subset of the peaks in promoter regions (**Figure S4C**).

**Fig. 4.**
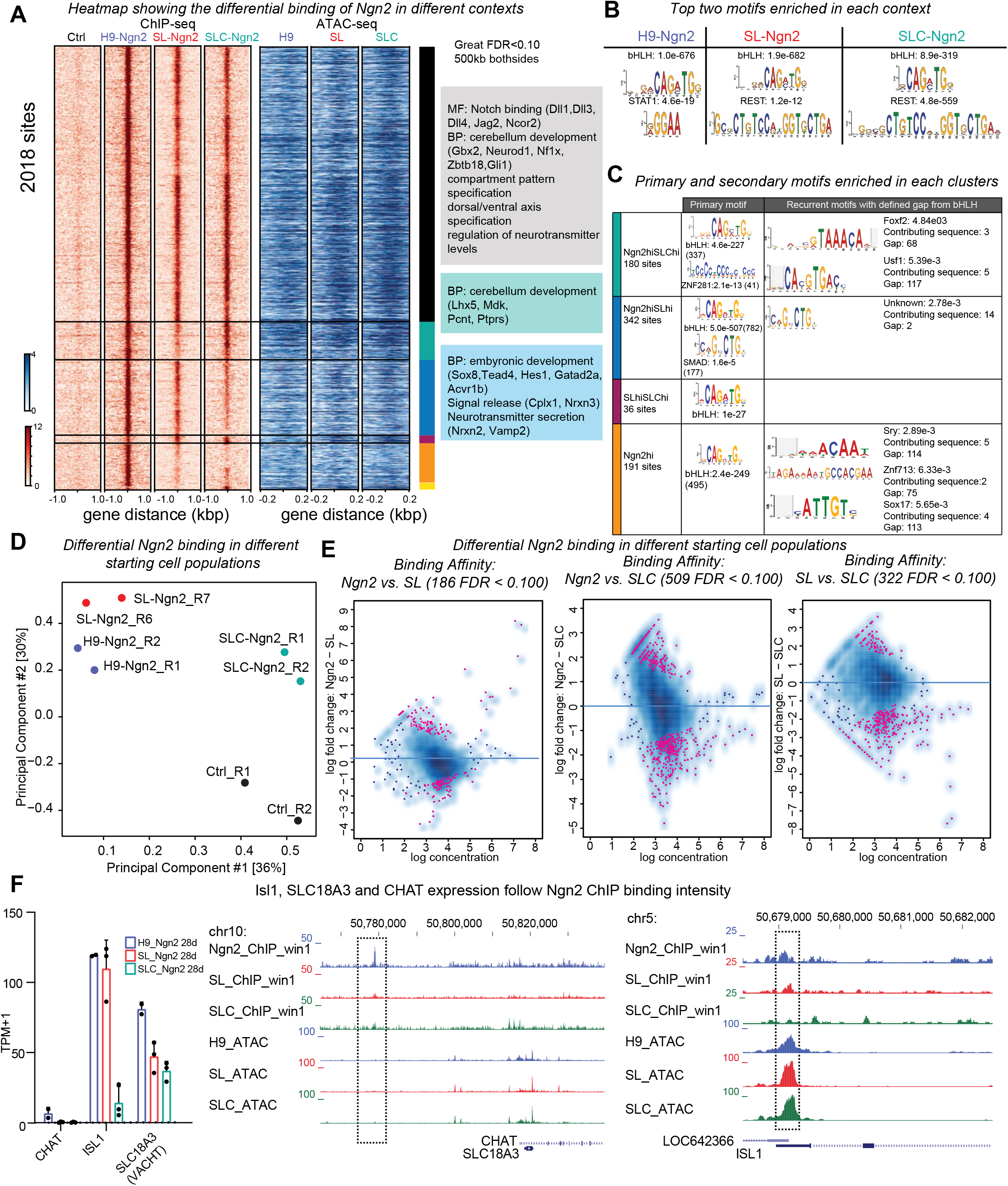
Genomic binding of NGN2 is chromatin dependent. (A) *Left (red heatmap)*: NGN2 ChIP sequencing profile two days after infection with Ngn2 in three different chromatin states (H9, SL, SLC) or with rtTA only infection (Ctrl) (n=2). Corresponding peaks are displayed +/− 1kb from the peak summit. *Right (blue heatmap)*: ATAC sequencing profile for the three starting populations (H9, SL, SLC) for the corresponding NGN2 ChIP-seq regions. The gene ontology terms were inferred using GREAT using genes within 500kb from the peaks (FDR p-value <0.10). (B) Top two motifs significantly enriched in the three populations (H9, SL, SLC) using sequences peaks called by MACS2 (C) Primary and secondary motifs with defined gaps from the primary motif significantly enriched in the sequences within +/− 250bp from the peak summit in four clusters (H9-NGN2^high^SLC-NGN2^high^, H9-NGN2^high^SL-NGN2^high^, SL-NGN2^high^SLC-NGN2^high^, H9-NGN2^high^) highlighted in Figure 4A. The number of sites with the motif was included in brackets. For each secondary motif, the E-value, the number of contributing sequences and the gap between the primary and the secondary motif are listed to the right. (D) Principal component analysis of the NGN2 ChIP in three different chromatin environments. The first principal component separates the different starting populations and the second principal component separates the control from the NGN2 treated three starting populations (E) Differential binding of NGN2 in three different chromatin environments analyzed by DiffBind (FDR p-value<0.10). There are 186, 506 and 322 differentially bound peaks when comparing NGN2 vs SL, NGN2 vs SLC and SL vs SLC, respectively. (F) *Left:* RNA-seq expression values (TPM+1) for the day 28 iN cells from the three starting populations (H9, SL and SLC) for *CHAT, ISL1 and SLC18A3*. Error bars = s.e.m. *Right:* Genomic tracks showing the NGN2 ChIP- and ATAC-seq signal at the *CHAT/SLC18A3* and *ISL1* loci. Note the correlation between the NGN2 ChIP peak heights and expression of corresponding genes shown on the left.

Given the surprising differential binding of NGN2 we next sought to better characterize these differences. Hierarchical clustering of NGN2 peak intensity (+/− 50bp from the peak summits) showed that in fact the majority of sites (about 60%) are commonly bound by NGN2 in all three cell types (**Figure 4A,** marked by black stripe). These common peaks are adjacent to genes that are enriched in GO terms such as *Notch binding* and *regulation of neurotransmitter levels*. The remaining 40% sites comprise 5 clusters that show preferential binding in one or two cell types. Notably, the majority of such sites comprising the 3 largest clusters show NGN2 binding in H9 cells with co-enrichment in SLC (**Figure 4A,** turquois cluster), co-enrichment in SL (**Figure 4A,** blue cluster), or no co-enrichment in either SL or SLC (**Figure 4A,** orange cluster). Only few sites are depleted in ES cells and enriched in both SL and SLC (**Figure 4A,** red cluster) or enriched in SLC only (**Figure 4A,** yellow cluster). Thus, unlike what we found for its closely related factor Ascl1, NGN2 genomic binding is clearly chromatin context dependent.

PCA analysis of the three ChIP-seq samples showed that H9-NGN2 and SL-NGN2 cluster more closely together than SLC-NGN2 (**Figure 4D**). When we performed differential NGN2 binding analysis among the three samples, we found that there 186, 509 and 322 peaks differentially occupied when comparing H9-NGN2 vs SL-NGN2, H9-NGN2 vs SLC-NGN2 and SL-NGN2 vs SLC-NGN2 (FDR <0.1) (**Figure 4E**). For example, NGN2 binds in all three conditions in the proximity of HES6 while only NGN2 in SLC-NGN2 condition bound to the distal region from the *LHX3* promoter (**Figure S4G, H**).

### NGN2 directly binds and regulates ISL1 and cholinergic genes

The prominent ISL1 expression in NGN2-iN cells was a surprising result, since NGN2 expression does not correlate well with Isl1 during development as only a small subset of NGN2 expressing progenitor cells give rise to Isl1+ neurons and not all cholinergic neurons are derived from NGN2+ cells. Moreover, we had found no correlation between the protein levels of NGN2 and ISL1 in human iN cells by immunofluorescence (**Figure S1E**). However we were surprised to find NGN2 bound at the *ISL1* and the combined *CHAT* and *vAChT (SLC18A3)* loci (**Figure 4F**, right traces). NGN2 binding strength correlated with the expression level of *ISL1*, *CHAT* and *vAChT (SLC18A3)* between the three cell types (**Figure 4F,** compare expression left with binding right).

### A combination of chromatin accessibility and signaling pathway-induced transcription factors may guide NGN2 binding

To characterize the chromatin state at differential NGN2 binding sites, we performed ATAC sequencing in the three populations (H9, SL and SLC) before and after NGN2 expression. Confirming the proper regionalization, the promoter region of the anterior gene *FOXG1* was most accessible in anterior (SL) cells and the posterior gene *CDX2* more accessible in posterior (SLC) cells (**Figure S4E**). Using PCA, we found that the neural SL and SLC cells cluster closer together than undifferentiated ES cells (**Figure S4D**). Following NGN2 expression all three populations shifted remarkably similarly into the same direction suggesting that the effects of NGN2 on chromatin is similar between the three cell types.

We next asked whether the differential chromatin accessibility may explain NGN2 binding. Indeed, in many cases, we found an overall correlation between NGN2 binding strength and the ATAC signal (**Figure 4A**). For instance, regions strongly bound in all three cell types are generally more accessible (*large parts of the black cluster*). In those clusters that are primarily bound in the neural SL or SLC cells but not ES cells (*yellow and red clusters*), the degree of chromatin accessibility correlated well with NGN2 binding. In those cases, the process of neuralization may have opened the chromatin configuration allowing NGN2 access.

However, other regions (*green and blue clusters*) cannot be explained by differential chromatin accessibility (**Figure 4A**) nor the types of E-boxes alone (**Figure S4F**). We thus performed motif enrichment analysis on these two clusters and found an additional enrichment of ZNF281 motif and SMAD4 motif adjacent to the bHLH motif in the H9-NGN2 and SL-NGN2 specific peaks (*blue*) and H9-NGN2 and SLC-NGN2 (*green*) respectively (**Figure 4C**). It was previously reported that both TCF and ZNF281 motifs are co-enriched in β-catenin ChIP-seq, suggesting ZNF281 might be downstream of Wnt signalling (Kjolby and Harland, 2017). This shows that the downstream effectors TGFβ/BMP and WNT (SMAD4 and ZNF281, respectively) may guide NGN2 to these differential sites even though they are in a less accessible state.

### EMX1 and FOXG1 cooperate with NGN2 to induce homogeneous forebrain excitatory neurons

In the mammalian brain, OTX2, OTX1, EMX2 and EMX1 show a nested and progressively restrictive expression pattern with OTX2 expression extending from the telencephalon to mid-hindbrain boundary and EMX1 limiting within the glutamatergic neurons, astroglia and oligodendrocyte of most pallial structures (Gorski et al., 2002; Yoshida et al., 1997) (**Figure 5A**). In our quest to further improve neuronal specification, we thus wanted to explore the effects of coexpressing these spatially restricted transcription factors together with NGN2. To this end, we cloned multiple forebrain and fore-/midbrain transcription factors (EMX1, EMX2, OTX1, OTX2, TBR2, LHX2 and FOXG1) and co-expressed them with NGN2 (**Figure 5B, S5A**) (Hébert and Fishell, 2008). All transcription factors, except OTX1 and TBR2, produced iN cells upon overexpression with NGN2 (**Figure S5A, S5C**). We first used ISL1 expression to assess whether the addition of forebrain transcription factors would focus the cellular subtype specification of NGN2-only iN cells. Indeed, when characterized on day 28 after infection, the additional expression of EMX1, EMX2 and FOXG1, but not OTX2, greatly reduced the percentage of ISL1-positive neurons (**Figure 5C, S5D**). In all conditions, the overwhelming majority of iN cells expressed the excitatory marker vGLUT (**Figure S5B, S5C**). We then performed RNA-sequencing. The PCA showed that NGN2:EMX1, NGN2:EMX2 and NGN2:FOXG1 are similar to each other and distinct from both NGN2 and NGN2 OTX2 (**Figure 5D**). Hierarchical clustering based on the genes that are significantly changed at least 2-fold within the different transcription factor combinations shows that both NGN2 with EMX1 and EMX2 overexpression induced genes that are involved in *cerebral cortex development* (*FOXP2*) and *anterior specification* (*BTG2, EMX2, NEUROD1*) and NGN2:OTX2 overexpression upregulated a large number of progenitor transcription factors (*NKX2-1, MEIS2, SOX6, NR2F1)* (**Figure 5E**). Furthermore, we found that all three cholinergic genes (*ChT (SLC5A7), vAChT (SLC18A3), and CHAT*) as well as *ISL1, PHOX2B* were repressed in NGN2:EMX1, NGN2:EMX2 and NGN2:FOXG1 compared to NGN2 or NGN2:OTX2 (**Figure 5F**). This result is confirmed in an independent cell line (**Figure S5H**). qRT-PCR for *ISL1, ChT (SLC5A7)* and *vGLUT1* confirmed the sequencing results (**Figure S5E**).

**Fig. 5.**
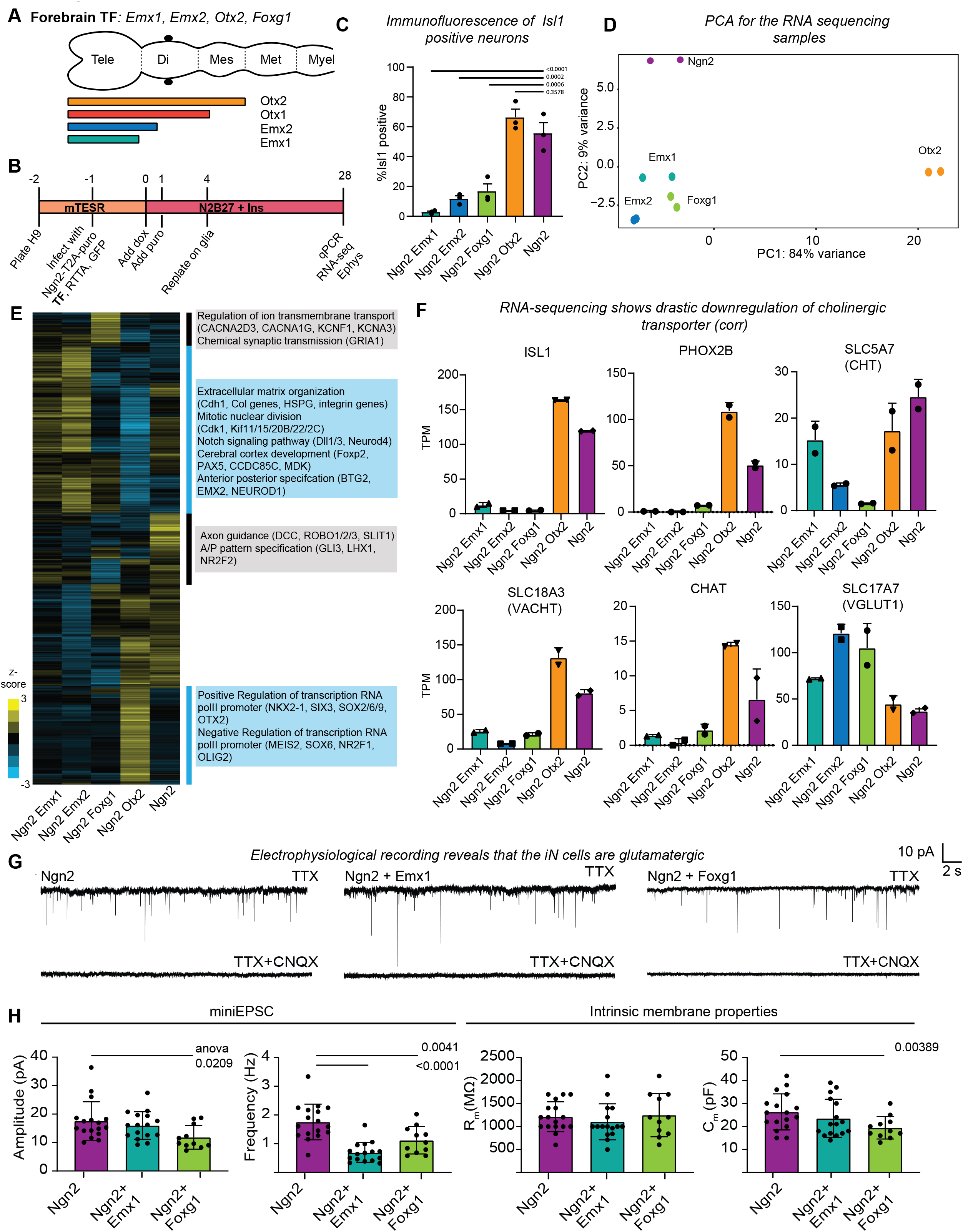
EMX1 and FOXG1 eliminate neurotransmitter subtype heterogeneity of Ngn2 iN cells. (A) Nested expression pattern of OTX2, OTX1, EMX2 and EMX1 during development (Tele=Telencephalon, Di=Diencephalon, Mes=Mesencephalon, Met = Metencephalon and Mye=Myelencephalon). (B) Protocol to generate Ngn2 iN cells co-expressing individual candidate forebrain transcription factors (TF). (C) Quantification of ISL1+ cells among all GFP/NGN2-infected iN cells on day 28. Representative fluorescence images are shown in Figure 5SD. (N=3, error bars = s.e.m. ANOVA statistical test was used. Exact adjusted p-values are shown above graph) (D) Principal component analysis of RNA-seq results of day 28 iN cells generated with NGN2 alone or in combination with EMX1, EMX2, FOXG1, or OTX2. (E) Hierarchical clustering of significantly changing genes in RNA-seq (>=2-fold change and p adj<0.05) within any two of the day 28 iN cells generated with NGN2, NGN2:EMX1, NGN2:EMX2, NGN2:FOXG1, or NGN2:OTX2. Significant gene ontology terms for the corresponding highlighted regions are listed. (P adj <0.05) (F) Bar graphs showing expression values of RNA-seq (TPM) for the two cholinergic transcription factors (ISL1, PHOX2B), three cholinergic genes (SLC18A3, SLC5A7, CHAT) and the glutamatergic marker (SLC17A7) of day 28 iN cells generated with NGN2 alone or in combination with EMX1, EMX2, FOXG1, or OTX2. (G) Representative traces of miniature excitatory postsynaptic currents (EPSCs) for day 28 NGN2-, NGN2:EMX1- and NGN2:FOXG1-iN cells in the presence of the voltage-gated Na+-channel blocker TTX to block action potential formation and network activity (top trace). All synaptic events observed are eliminated after addition of the excitatory AMPA receptor inhibitor CNQX (bottom trace). (H) Quantification of miniature EPSC amplitude and frequency and the two intrinsic membrane properties input resistance and capacitance of NGN2, NGN2:EMX1, and NGN2:FOXG1 iN cells. (P-values using the ANOVA test are shown above bars. N=17, 16, 11 cells measured in 3 independent batches in NGN2, NGN2:EMX1 and NGN2:FOXG1 28d iN cells plated on glia. Error bars = s.e.m.)

Similarly, inspecting other neurotransmitter subtype specific genes, EMX1, EMX2, and FOXG1 repressed most monoaminergic subtype specific genes while maintaining the level of glutamatergic subtype specific genes (**Figure S5I**). All three genes critical for GABAergic identity were even more reduced in NGN2 only cells by co-expression of EMX1 and 2. FOXG1 reduced expression of both Glutamate decarboxylases 1&2 but left vesicular GABA transporter (vGAT/SLC32A1) unchanged (**Figure S5I**). Of note, EMX1 is only expressed in excitatory neurons whereas FOXG1 is expressed in both excitatory and inhibitory neurons (**Figure S5F**).

These data show that in particular EMX1 potently eliminates the cholinergic, monoaminergic, and GABAergic programs partially induced by NGN2 alone resulting in much more homogeneous excitatory cells. We therefore wondered whether the failure to eliminate cholinergic program by anterior patterning (SL vs. SLC cells) that we had observed before could perhaps be explained by a lack of *EMX1* expression. Indeed, while *EMX1* was properly induced in anterior progenitor cells, it was rapidly downregulated during neuronal differentiation and we found an anti-correlation between *EMX1* and ISL1 expression levels in day 2 and 28 cells (**Figure S5G**).

Finally, we sought to functionally characterize the newly generated excitatory cells. While EMX1, EMX2, OTX2 and FOXG1 are well-characterized developmental regulators, their expression in adult human cortical excitatory neurons is less known. Analyzing data from the Allen brain atlas showed that OTX2 is not expressed well in the adult cerebral cortex but FOXG1 is prominently expressed throughout all neural cell types except oligodendrocytes, EMX1 is quite restricted to excitatory neurons and EMX2 is most strongly expressed in astrocytes (**Figure S5F**). Thus, among those 4 genes, only EMX1 and FOXG1 are expressed in cortical excitatory neurons. We therefore tested whether NGN2-iN cells with and without EMX1 and FOXG1 co-expression are glutamatergic by electrophysiology. All iN cells analyzed from the three groups exhibited miniature excitatory postsynaptic currents (EPSCs) that could be blocked by the application of an AMPA receptor antagonist, CNQX (**Figure 5G**). Of note, upon AMPA receptor blockade no other kinds of postsynaptic currents were detectable showing that these iN cells are predominantly glutamatergic. The EPSC amplitudes were similar between NGN2 and NGN2:EMX1 and slightly decreased in NGN2:FOXG1 iN cells (**Figure 5H**). The miniEPSC frequency was slightly decreased in NGN2:EMX1 and NGN2:FOXG1 when compared to NGN2 cells (**Figure 5H**). The intrinsic membrane properties such as membrane resistance and capacitance, parameters influenced by overall cell size and ion membrane permeability, were similar between the different conditions (**Figure 5H**).

### The forebrain transcription factors EMX1 and FOXG1 redirect NGN2 chromatin binding explaining decreased activation of cholinergic genes

Next, we wanted to explore the mechanisms how the addition of EMX1 or FOXG1 restrict NGN2’s ability to induce neuronal subtypes. There are two non-mutually exclusive principle possibilities: (i) EMX1/FOXG1 act independently of NGN2 and influence gene expression in an additive manner or (ii) they could directly influence the targeting of NGN2 to the chromatin. To distinguish between these two possibilities, we expressed a FLAG-tagged version of NGN2 in hES cells and performed FLAG antibody ChIP-sequencing with and without co-expression of EMX1 and FOXG1 (**Figure 6A**). Remarkably, the NGN2 binding patterns were distinct between the three conditions with some shared and some condition-specific binding sites (**Figure 6A**). Differential NGN2 binding analysis revealed 1603, 2701 and 1525 differential peaks in pair-wise comparisons of the three conditions (FDR <0.1) (**Figure 6B**). For example, the distal region of the FOXO6 locus is bound by NGN2 in ES cells only when co-infected with FOXG1 (**Figure S6D,** green highlight) whereas the NGN2 binding site at the HES6 locus is bound in all conditions (**Figure S6E**, purple highlight). The FOXG1- and EMX1-induced modulation of NGN2 binding is also reflected in the PCA analysis that shows a clear separation primarily by the second principal component (**Figure S6F**). When compared to differential regionalization of neural precursor cells (SL and SLC cells), EMX1 or FOXG1 co-expression led to a much more enhanced relocation of NGN2 in ES cells (compare **Figures 6B and 4E**).

**Fig. 6.**
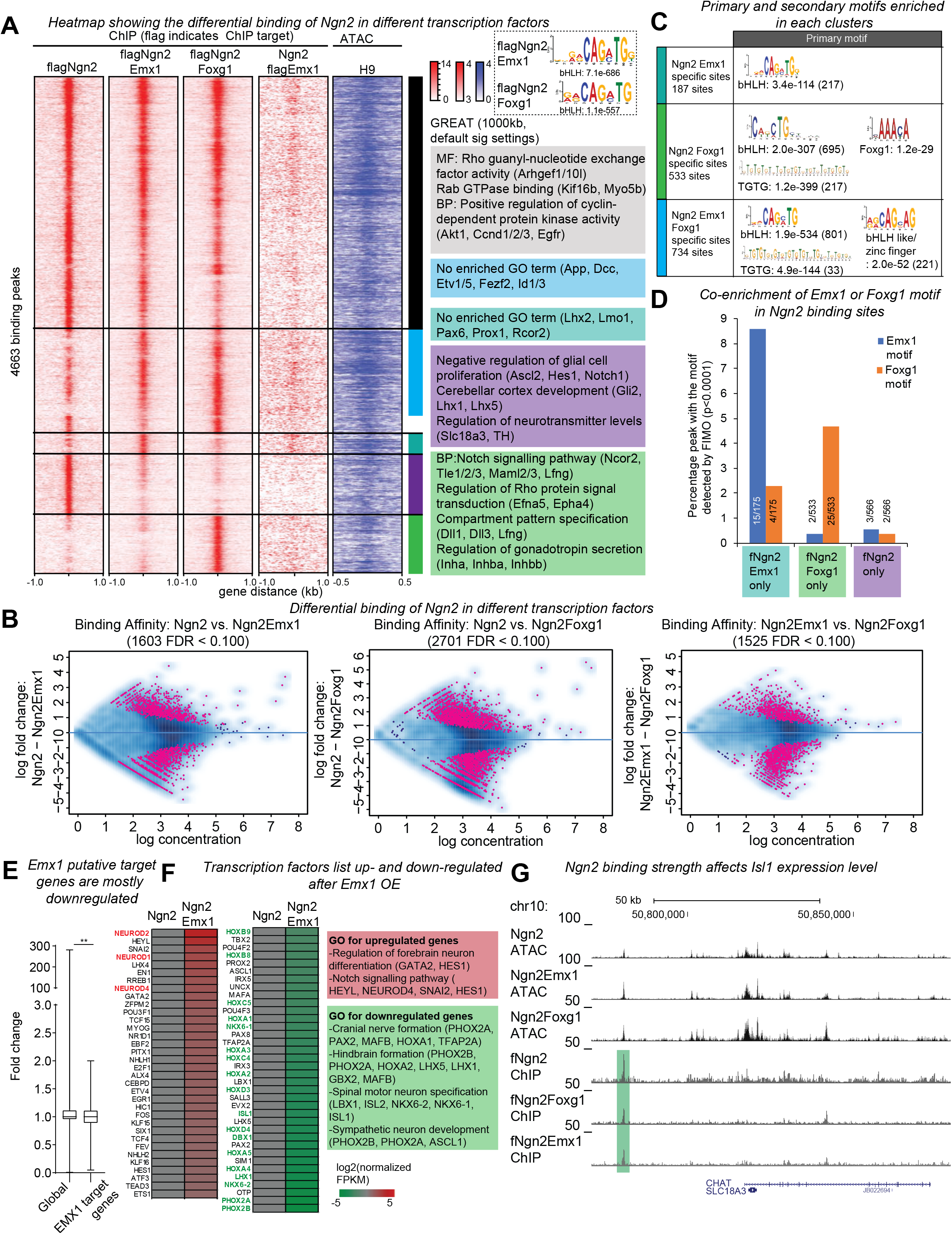
EMX1 and FOXG1 change NGN2 chromatin targeting. (A) *Left three columns*: flagNGN2 ChIP-seq profile within +/− 1kb from the peak summit (red signal) in ES cells two days after infection with flagNGN2 alone or with EMX1 or FOXG1 (n=2). *4^th^ column*: Corresponding genomic regions as in left 3 columns showing the flagEMX1 ChIP-seq profile 2 days after infection with flagEMX1 and NGN2. All ChIP-seq peaks are displayed +/− 1kb from the peak summit. *5*^*th*^ *column (blue signal)*: Corresponding genomic regions showing ATAC-seq signal in ES cells (H9). The ATAC-seq peaks are displayed +/− 500bp from the respective flagNGN2 ChIP peak summit. Gene ontology terms for various genomic clusters called significant by GREAT analysis are shown using genes within 1000kb from the peaks. Blue and turquois clusters did not significantly enrich any GO term. *Dotted box:* Position weight matrix of the bHLH motif enriched by de novo motif search analysis of the flagNGN2 ChIP-seq of flagNGN2:EMX1 and flagNGN2:FOXG1 infected cells considering all significant peaks (+/−50bp from the peak summit) (B) Pairwise comparison of differential NGN2 binding in three different conditions (flagNGN2, flagNGN2:EMX1, flagNGN2:FOXG1 infection) analyzed by DiffBind (FDR p-value<0.10). There are 1603, 2701 and 1525 differentially bound peaks when comparing flagNGN2 vs flagNGN2:EMX1, flagNGN2 vs flagNGN2:FOXG1 and flagNGN2:EMX1 vs flagNGN2:FOXG1. (C) The top motif significantly enriched by motif search analysis of three clusters color-indicated in A: NGN2 peaks specific to NGN2:EMX1 infected cells (turquois cluster); NGN2 peaks specific to NGN2:FOXG1 infected cells (green cluster); NGN2 peaks specific to NGN2:EMX1 and NGN2:FOXG1 infected cells (blue cluster). Sequences +/− 500bp from the peak summit were used. The number of sites with the motif was included in brackets. (D) Reverse motif search for EMX1 and FOXG1 motifs in three different clusters showing differential NGN2 binding: NGN2 peaks unique to cells infected with flagNGN2 and EMX1 (turquois cluster); NGN2 peaks unique to cells infected with flagNGN2 and FOXG1 (green cluster); and NGN2 peaks unique to cells infected with flagNGN2 alone (purple cluster). Shown are percentages of motifs among total number of genomic sites in the respective cluster. Numbers in bars show the actual numbers. The positional weight matrix used for EMX1 and FOXG1 motifs were obtained from Jaspar. (E) Boxplot showing the expression fold change of all genes and predicted Emx1 target genes by RNA-seq in NGN2 vs. NGN:EMX1 infected cells. Plotted are TPM+1_NGN2:EMX1_/TPM+1_NGN2_ of all genes and genes within 10kb of an EMX1 peak. Predicted EMX1 target genes are significantly repressed (average fold change of all genes 1.01; average fold change of predicted EMX1 targets 0.89; p value = 0.0027) (F) Heatmap showing transcription factors regulated by EMX1 in the context of NGN2 expression. Boxes to the right show the GO terms enriched in either group and the genes included in the GO terms. (G) Genomic tracks showing the ATAC and NGN2 binding peaks at the *CHAT/SLC18A3* locus in different conditions. Note the NGN2 peak height in the three different conditions (green highlight) corresponds to the gene expression levels shown in Figure 5F.

We next wondered whether the altered NGN2 binding pattern may explain the prominent repression of cholinergic genes. Indeed, a prominent NGN2 peak upstream of the *CHAT/vAChT (SLC18A3)* locus was weaker bound by NGN2 when either FOXG1 or EMX1 were co-expressed (**Figure 6G**). Then, we wanted to find potential explanations for the differential NGN2 binding. Sites that are unique in NGN2:FOXG1 cells were enriched for *regulation of gonadotropin secretion* and *compartment pattern specification* consistent with the FOXG1 expression pattern (**Figure 6A**). Sites unique for the NGN2-only condition enriched for the GO terms *negative regulation of glial cell proliferation, regulation of neurotransmitter levels,* and *cerebellar cortex development*. No GO terms were significantly enriched in sites unique to NGN2:EMX1 (**Figure 6A**).

To understand whether differences in binding among the groups can be explained by chromatin accessibility alone, we plotted the ATAC sequencing signal in hES cells centering at the flagNGN2 peak summits and found that the different clusters are all similarly accessible (**Figure 6A,** blue). Thus, the unique NGN2 binding sites in NGN2:EMX1 and NGN2:FOXG1 are not simply due to inaccessibility in ES cells.

We next performed motif and peak distribution analysis of the NGN2 peaks. In all NGN2 ChIP-seq datasets a very similar bHLH motif (CA<u>GA</u>TG) was enriched and most NGN2 peaks harbored at least one bHLH motif and often multiple (**Figure 6A, S6A**). When we further subdivided bHLH motifs enriched in the NGN2:EMX1 and NGN2:FOXG1, we found that NGN2:EMX1 has a higher percentage of the CA**GC**TG E-box motif and NGN2:FOXG1 has a higher percentage of the CA**GA**TC E-box motif similar to NGN2 alone (**Figure 6C, S6B**). The peak distribution was similar between the samples (**Figure S4C, S6C**).

The differential E-Box enrichment among different NGN2 peaks suggested potentially additional sequence similarities among the different NGN2 bound clusters. Indeed, the unbiased de-novo motif search analysis also produced the canonical Foxg1 motif among the NGN2:FOXG1-unique NGN2 binding sites (**Figure 6C**). Reverse motif search showed that the Emx1 motif was four times more enriched under the NGN2 peaks specific in the NGN2:EMX1 cells and the Foxg1 motif was eight times more enriched within the NGN2 peaks specific for the NGN2:FOXG1 cell compared to the peaks specific for the NGN2-only infected cells (**Figure 6D**). Remarkably, the Emx1 motif was also more enriched in NGN2:FOXG1 specific peaks and the Foxg1 motif in NGN2:EMX1 specific peaks. Even though to date EMX1 or FOXG1 are not known to physically interact with NGN2, these data suggest that EMX1 and FOXG1 may recruit NGN2 to their own binding sites, even when they contain less preferred E-box motifs.

To explore this idea further, we performed ChIP-sequencing for flagEMX1 in ES cells co-infected with flagEMX1 and NGN2. We obtained 1393 significant peaks (n=2, idr<0.10) (**Figure S6G**) and most peaks contained a homeobox motif (**Figure S6G, I**). They are located in the intergenic, intron and gene body (**Figure S6H, J**). The peak distribution of EMX1 was slightly different than NGN2 with a much reduced promoter region localization (**Figure S6J,** compare to **Figure S6C**). When plotting the flagEMX1 ChIP-seq signal from genomic sites that are bound by NGN2, we could indeed see an enriched EMX1 binding at NGN2 peaks unique in the NGN2:EMX1 infected cells (dark green cluster in **Figure 6A**). This demonstrates that cooperative binding with EMX1 allows NGN2 to bind new genomic sites.

### EMX1 represses posterior and cholinergic genes independent of NGN2

We next explored whether EMX1 may have NGN2-independent functions, in addition to directly influencing NGN2 chromatin binding. In particular we were wondering whether EMX1 may repress the promiscuous activation of neuronal subtypes programs. Therefore, we first explored the direct transcriptional effects of EMX1. We identified the putative EMX1 target genes (genes within 10Kb of EMX1 binding) and plotted their average expression level in ES cells infected with NGN2 or EMX1:NGN2. We found a lower average fold change and narrower distribution of fold change of these EMX1 target genes when the cells where coinfected with EMX1 compared to NGN2 alone (**Figure 6E**). This indicates that EMX1 might have repressive functions. Repressed genes include posterior genes like HOX genes, ISL1, and PHOX2 genes, involved in cranial nerve, hindbrain formation and spinal and sympathetic neuron development. On the other hand anterior genes involved in forebrain development were upregulated upon EMX1 co-expression (**Figure 6F**). In support of this notion, we found that Emx1 binds the *ISL1* locus when expressed in ES cells (**Figure S6H**). Thus, EMX1 acts independently of NGN2 to repress posterior genes but also influences NGN2 binding to achieve a homogeneous forebrain glutamatergic neuronal identity.

## DISCUSSION

In this study we used NGN2 hES-iN as a platform to systematically investigate the effects of epigenetic landscapes induced by differentiation and the effects of introducing transcription factors with a spatially restricted expression pattern. During development, neural progenitor cells acquire a regional identity via spatially restricted transcription factors induced by morphogens. The proneural bHLH transcription factor, NGN2 then “primes” the neural progenitor cells with pan-neuronal genes and a neurotransmitter subtype specific gene program.

When expressed in human ES cells, NGN2 induces neurons of exclusively glutamatergic subtype specification. However, our single cell characterization here revealed a certain degree of heterogeneity of NGN2-iN cells with partial expression of cholinergic and monoaminergic programs among a subgroup of excitatory cells. We found that unexpectedly, cholinergic programs are directly induced by NGN2 in human ES cells, via direct regulation of cholinergic effector genes and the developmental regulator ISL1. In the absence of a step to specify regional identity, the incomplete neurotransmitter programs were retained in addition to the glutamatergic program in neurons. Therefore, we deduced that adding a step to endow the cells with a regional identity by differentiation would eliminate the unwanted cholinergic program. We chose to differentiate human ES cells into anterior and posterior neuroectodermal cells before NGN2 induction. NGN2 is highly expressed in glutamatergic cortical projection neurons (anterior) and in the cholinergic spinal motor neurons (posterior) (Parras et al., 2002). In those experiments, we showed that the incomplete cholinergic and monoaminergic programs can be modified but not eliminated with a more defined starting population. For example, iN cells obtained from the anterior neural stem cells have a higher level of vGluT1 (SLC17A7) (a glutamatergic marker) and a low level of ChT (SLC5A7) (a cholinergic marker) when compared to the posterior neural stem cells, fitting the neuronal population NGN2 is expressed (**Figure 3I**). However, not all cholinergic genes were completely downregulated. iN cells derived from the anterior and posterior neural stem cells have similar expression levels of two cholinergic genes, vAChT (SLC18A3) and CHAT. This could be due to the two starting populations we used. The anterior neural stem cells we used in this paper express *OTX2* whose expression span telencephalon, diencephalon and mesencephalon (**Figure S4I**). Thus, we cannot rule out further refinements in the differentiation protocol (instead of only anterior and posterior neural cells) may affect the neurotransmitter identity and further specify a more homogeneous neuronal subtype.

Unlike neurotransmitter phenotype specification, we found that the regional identity of mature iN cells is highly dependent on that of the starting population. This finding is significant as it opens up the NGN2 iN cell platform to generate glutamatergic neurons with different regional identities by merely changing the starting population. More importantly, the regional identity of the starting population instructs which sets of genes are upregulated or downregulated (**Figure 3F, G**). We think that different regional identities of the starting population might present NGN2 with different chromatin landscapes, resulting in different lineage specification genes being regulated. Indeed, we found regions that are differentially bound by NGN2 among the three starting cell populations (**Figure S4I**). A subset of the differentially bound regions can be attributed to chromatin accessibility alone. The other differentially bound regions could be explained by the secondary motifs found which we attributed to the downstream effectors of TGFβ/BMP and WNT pathways. Moreover, even for the commonly bound sites, the peak height might be different. We found that the NGN2 peak height correlates with the expression of the closest genes. Given that peak height is a proxy to estimating the proportional of the cells having the peak and that NGN2 binds mostly to enhancer regions, chromosome conformation assays could be used in the future to examine this hypothesis. Summing up, we have shown NGN2 binds differently in different starting populations, inducing vastly different downstream genes.

We next asked whether region-specific transcription factors can modulate NGN2-induced neuronal specification. We shortlisted a handful of candidates with varying degree of restrictive expression in telencephalon and co-expressed them individually with NGN2(Hébert and Fishell, 2008). EMX1 and FOXG1 gave the clearest results when overexpressed with NGN2. We observed a precipitous decrease in the level of transcription factors involved in cholinergic neuron specification (*ISL1* and *PHOX2B*) and all cholinergic rate limiting enzyme and vesicular proteins (*CHAT, ChT (SLC5A7), vAChT (SLC18A3)*). During development, *EMX1* expression starts in mouse around E9.5 and is expressed in differentiating and mature mostly glutamatergic neurons and astrocytes in the cortex, hippocampus and olfactory bulb (Briata et al., 1996; Yoshida et al., 1997). *FOXG1* expression begins around E8.5 in the telencephalic primordium which later becomes dorsal and ventral telencephalon (Tao and Lai, 1992). Dorsal telencephalon includes neocortex and hippocampus and generates predominantly glutamatergic neurons. Ventral telencephalon includes medial, lateral and caudal ganglionic eminences and generates various populations of mostly GABAergic neurons in the cortex, striatum, basal ganglia and olfactory bulb. It is thus not surprising to see an additional incomplete GABAergic program, indicated by the SLC32A1 (or VGAT) expression, in the NGN2 FOXG1 day 28 iN cells. In the same note, OTX2, albeit lowly expressed in mature cortical excitatory and inhibitory neurons, induced additional cholinergic and dopaminergic programs when co-overexpressed with NGN2. OTX2 is believed to be a selector type transcription factor (Arlotta and Hobert, 2015) as conditional deletion OTX2 in the neural stem cells using Nestin-cre led to the formation of cerebellar-like structure instead of the colliculi and ectopic formation of serotonergic neurons in the midbrain (Vernay et al., 2005). All these data support the notion that NGN2, when restricted by other homeodomain transcription factors with distinct A-P expression domain, upregulates neurotransmitter programs of the neurons later present in that domain and downregulates unwanted neurotransmitter programs (**Figure S6K**). The effects of the homeodomain transcription factors *in vitro* reported here are in agreement with the role of “spatial selectors” during development which are defined as factors with defined spatial expression that instruct/restrict the regional identity of multipotent progenitors and later limit the subtypes of mature neurons generated from that progenitor domain (Allan and Thor, 2015). This function is similar to the “many-but-one” cell type specification function of MYT1L. Just like MYT1L limits the programs of many other lineages except the neuronal, EMX1 represses all other neurotransmitter subtype programs except the glutamatergic program (Mall et al., 2017). Given that *EMX1* and *FOXG1* expression are also detected in adult forebrain projection neurons as reported in scRNA-sequencing data, we reckon that in an *in vitro* setting EMX1 and FOXG1 might also have terminal selector functions where it actively promotes the correct neurotransmitter subtype genes and limits the promiscuous neurotransmitter subtype genes in mature iN cells from NGN2 protocol (**Figure 6F, S6K**) (Arlotta and Hobert, 2015; Hobert, 2008). In summary, these spatially defined homeodomain transcription factors present in neural progenitors thus restrict the promiscuity of NGN2 binding and assist in the generation of different neurotransmitter subtypes from a limited number of bHLH proneural transcription factors.

In this paper, we outlined the transcriptional neuronal input-output codes by modulating the chromatin environment of the starting populations or by pairing NGN2 with homeodomain transcription factors which are expressed in the progenitors and mature neurons. This protocol coupled with pre-differentiated neuronal progenitors using established protocols followed by overexpression of proneuronal factors or subsequent expression of subtype specific POU or homeodomain transcription factors would vastly generalize the utility of NGN2 iN cell protocols (Fattahi et al., 2016; Tchieu et al., 2017; Tsunemoto et al., 2018). This paper serves as a blueprint to generate not only bona fide neurons of different neurotransmitter subtypes and neurons of the same neurotransmitter subtypes residing in different anterior-posterior levels but also various cell types in the other organ systems (Guo and Morris, 2017).

## AUTHOR CONTRIBUTIONS

C.E.A and M.W. conceived and designed the experiments. C.E.A., V.O., B.Z., Q.Y.L, M.M., A.N., K.C., performed the experiments and analyzed the data. C.E.A and M.W. wrote the paper with input from all authors.

## ACKNOWLEDGEMENTS

This study was supported by a training grant from a training grant from California Institute of Regenerative Medicine (CIRM, TGR-01159), Siebel Foundation and Stanford Graduate Fellowship to C.E.A. Q.Y.L. was supported by the Singapore Agency for Science, Technology and Research (A*STAR). This project was also supported by a New York Stem Cell Foundation Robertson award (to M.W.), a Howard Hughes Medical Institute Faculty Scholar award (to M.W.), and a Tashia and John Morgridge Faculty Scholar award (to M.W.) by the Child Health Research Institute (CHRI) at Stanford. We thank members of the Wernig lab and Kyle Loh for insightful discussions. The sequencing in the paper was performed with help from Stanford Functional Genomic Facility and Stanford PAN facility.

## MATERIALS AND METHODS

### Reprogramming of human embryonic stem cells to induced neurons (iN)

We followed the protocol previously described (Zhang et al., 2013). Human embryonic stem cells (H1 and H9, University of Wisconsin) were plated single cell and infected the next day with TetO-FLAG-NGN2-T2A-PURO^R^ and FUW-rtTA. Doxycycline was added to the wells the next day. To select for only NGN2 transducing cells, puromycin (Final concentration: 2μg/ml, Sigma) was added in addition to doxycycline the next day and kept for 3 days. For prolonged culture, the cells were dissociated using Accutase and replated on mouse glia at day 4. For single cell RNA sequencing, doxycycline was added to 14d and removed for the last 14d.

For the knock-down experiment, shRNAs obtained from Sigma (ISL1: TRCN0000014893, TRCN0000014897; PHOX2B: TRCN0000358499, TRCN0000358500) were packaged into lentiviruses and co-infected with TetO-FLAG-NGN2-T2A-BLAST^R^ and FUW-rtTA. Doxycycline was added to the wells the next day. To select for only NGN2 transducing cells, puromycin (Final concentration: 2μg/ml, Sigma) and blasticidin (Final Concentration: 10μg/ml, Sigma) was added in addition to doxycycline the next day and kept for 3 days

For the forebrain transcription factors experiments (EMX1, EMX2, FOXG1, OTX2), they were first cloned into TetO-IRES-HYGRO^R^ plasmid and coinfected with TetO-FLAG-NGN2-T2A-PURO^R^ and FUW-rtTA. In the case of forebrain transcription factor experiment, hygromycin (150μg/ml, Roche) was added to select for the additional transcription factor.

For all NGN2 ChIP-sequencing experiment (NGN2 in different chromatin environments and NGN2 with EMX1/FOXG1), FUW-rtTA and TetO-FLAG-NGN2-T2A-PURO^R^ were used. For the EMX1 ChIP-sequencing experiment, FUW-rtTA, pTight-NGN2-PGK-puro and TetO-flagEMX1-IRES-HYGRO^R^ were used instead. The cells were dox induced for 2d before they were harvested and used for ChIP-seuqencing

The experiments were performed in accordance with California State Regulations, CIRM Regulations and Stanford's Policy on Human Embryonic Stem Cell Research.

### Differentiation of human embryonic stem cells into human anterior and posterior neural stem cells

We followed the protocol previously described (Du et al., 2015). Human embryonic stem cells were plated in clumps in the presence of Y27632 (10mg/ml, 1000x, Axon MedChem). The hES colonies were allowed to grow in mTESR for another day before the media was switch to differentiation media (1X N2, 1X B27, DMEM/F12: Neurobasal=1:1 (Invitrogen), 0.1mM Ascorbic acid (Sigma)) with small molecules [SB431542, LDN193189 and CHIR99021 (Final concentration: 10μM, 100nM and 3μM from Stemgent and Tocris)] and the cells were allowed to differentiate in the media for 6 days before they were dissociated with Accutase and plated at the density of 1 million cells per well of 6 wells for infection.

### Single cell RNA sequencing library preparation

We followed the Smart-seq2 protocol outlined (Picelli et al., 2013). iN cells at 4d and 28d were sorted in fluorescence activated cell sorter (BD FACS Aria) in the single cell mode into 96 well plates containing lysis buffer. The plates were snap frozen in dry ice for future processing. Poly(A)+ mRNA was enriched using an oligo(dT) primer in the first strand synthesis. PCR amplified cDNA for each cell was examined using fragment analyzer (Advanced Analytical) and the concentration was normalized before it was used for library preparation (Nextera XT kit, Illumina). Primers were removed using AMpure XP beads (Beckman Coulter) and the resulting library was sequenced paired ends and 150bp in the HiSeq 4000 platform.

### Single cell RNA sequencing analysis

Raw reads were processed as follow: adapter sequences were trimmed, and poor-quality score were removed using Skewer using the following settings (-q 21 -l 21 -n -u). Groomed reads were mapped to hg19 using STAR in a 2-pass setting (https://github.com/alexdobin/STAR). The FPKM values were obtained using Cufflinks [Cuffquant: using default settings and a hg19 gtf file, Cuffnorm: default settings]. Cells with <400000 reads were excluded from subsequent analysis. PCA and tSNE analysis were performed in R using FactorMineR and Seurat (y-cutoff=0.75, x-cutoff=0.5) respectively. Violin plots were drawn using ggplot2 in R.

### Bulk RNA sequencing

Total RNA_was collected using Trizol (Invitrogen) followed by cleanup using RNA Clean and Concentrator (Zymo) using the manufacturer’s protocol. Samples were then QCed using bioanalyzer and subjected to paired end sequencing (BGI platform).

### Bulk RNA sequencing analysis

Raw reads were mapped to hg19 and counts were obtained using STAR with the following settings (--runThreadN 16 --outSJfilterReads Unique --outFilterType BySJout --outSAMunmapped Within -- outSAMattributes NH HI AS NM MD --outFilterMultimapNmax 20 --outFilterMismatchNmax 999 --alignIntronMin 20 -- outFilterMismatchNoverReadLmax 0.04 --alignIntronMax 1000000 --alignSJoverhangMin 8 --alignSJDBoverhangMin 1 --sjdbScore 1 --outSAMtype BAM SortedByCoordinate --outBAMcompression 10 --limitBAMsortRAM 60000000000 --quantMode TranscriptomeSAM GeneCounts --quantTranscriptomeBAMcompression 10 ‒outSAMstrandField intronMotif) (Dobin et al., 2012). TPM were obtained from the mapped by using RSEM with the following settings (--paired-end --seed-length 21 --no-bam-output --calc-pme --calc-ci ‒alignments). Differential analysis was performed using DESeq2 in default settings (Love et al., 2014). Gene ontology analysis was performed using PANTHER and only GO terms that are significant (corrected p-value) were listed.

### ATAC sequencing analysis

Libraries were made following the protocol by Buenrostro and coworkers (Buenrostro et al., 2013) and sequenced paired ended in the Next-seq platform. Reads were trimmed using in-house scripts and mapped to the reference genome (hg19) using bowtie2 using the following additional option (--very-sensitive). Mitochondrial reads were then removed (awk ‘\$3!=“chrM”’). Reads were subsequently deduplicated using Picard (MarkDuplicates.jar). Mapped reads were examined to ensure enrichment in the transcription start sites and gave a characteristic fragment size distribution as described in the paper above.

### Chromatin immunoprecipitation followed by sequencing (ChIP-seq)

NGN2 with a N-terminal FLAG-tag was used. hES or anterior/posterior neural stem cell was infected with freshly purified viruses (TetO-FLAG-NGN2-T2A-PURO and FUW-rtTA) at day 0. After 48 hours post doxycycline induction, cells were dissociated with Accutase, crosslinked in 1% formaldehyde (Thermo Scientific) for 10min, quenched with 0.125mM glycine (Sigma, final concentration) for 5min and snap frozen in liquid nitrogen. 60-100 million cells were used per replicate. Flag and mouse IgG beads (Sigma) were washed with IP dilution buffer (1% Triton X-100, 2mM EDTA pH8.0, 20mM Tris-HCl pH8.0, 150mM NaCl, 1mM DTT, 100uM PMSF), blocked overnight at 4°C in IP dilution buffer with 0.1% BSA and 0.06% sheared salmon sperm DNA before use.

Nuclei were isolated by incubating with cell lysis buffer (5mM HEPES pH7.9, 85mM KCl, 0.5% NP40, 100uM PMSF, protease inhibitors from Roche) for 10min on ice and centrifuged at 5000rpm for 5min at 4°C, then lysed with nuclear lysis buffer (50mM Tris-HCl pH8.0, 10mM EDTA pH8.0, 1% SDS, 100uM PMSF, protease inhibitors) for 10min on ice. Chromatin was sheared using Covaris sonicator until DNA was fragmented to 200-500bp (Settings: 10 min, duty cycle =5%, intensity=4, cycle/burst=200). Sheared chromatin was diluted using 3X volume of IP dilution buffer and pre-cleared for at least 4hr at 4°C using IgG beads (Sigma A0919). 1% of pre-cleared chromatin was kept as input, and the remainder was incubated with FLAG beads overnight at 4°C. Beads were washed 8 times with IP wash buffer (20mM Tris-HCl pH8.0, 2mM EDTA pH8.0, 250mM NaCl, 1% NP40, 0.05% SDS, 100uM PMSF) and once with TE buffer with 100uM PMSF. Beads and 1% input samples were reverse cross-linked overnight in IP elution buffer (50mM NaHCO3, 1% SDS) at 65°C.

### ChIP-seq sequence alignment and peak calling

Reads were aligned to the hg19 reference sequence using Bowtie 2.1.0 (Langmead and Salzberg, 2012) with an additional −5 10 parameter to remove the low quality initial read. Peak calling was performed using MACS 2.1.1 (Zhang et al., 2008). For experiments with two replicates, reproducible peaks were selected using IDR 2.0.2 (Li et al., 2011) with cutoff of idr ≤ 0.1 using the recommended pipeline for MACS2. For experiments with more than 3 replicates, pairwise IDR was performed and reproducible peaks (idr ≤ 0.1) for each pair were merged into a single peak list using Bedtools merge. Differential peak binding and principal component analysis were performed using Diffbind with summit=50 at q<0.10(Ross-Innes et al., 2012). Peaks were considered differential bound if they have q<0.10 by DiffBind. The profile of peaks that are differently expressed were then plotted using deeptools suite following the vignette. Motif enrichment analyses were performed in the MEME-ChIP online suite using default settings with the following changes (MEME options: Any number of repetitions, No of motifs should MEME find: 10). Gene ontologies for the genes cis to the ChIP peaks were performed using GREAT.

### Western blotting

In order to measure the protein levels 10μl of sample buffer was added for every 50000 cells and 1μl of benzonase was added to the samples. The samples were placed on a heat block (55°C) for 10-15 minutes. A precast 8-12% bis-tris gel was then loaded into the chamber, filled with MES buffer containing 50 mL of running buffer (Invitrogen) and 950 mL of deionized H_2_O. 10μl of the denatured sample and a protein ladder was added per well. The gel was run at 200V and then transferred onto a PVDF membrane by running the transfer at 40V for 2 hours on ice in a transfer buffer (850 mL deionized H2O, 100 mL methanol, and 50 mL of transfer buffer) (Invitrogen). The membrane was then blocked with 5% milk in phosphate buffer saline with 0.1% Tween-20 for 30 minutes and stained with a primary antibody overnight. The gel was then stained with a secondary antibody, imaged, and scanned for analysis. Antibodies used: goat-OTX2 (1μg/ml. R&D), rabbit-HSP90 (1:2500, Cell Signaling), rabbit-FLAG (1:500, Sigma), mouse-FLAG (1:1000, Sigma), mouse-ChT (1:500, Synaptic System). Goat and rabbit conjugated horseradish peroxidase (1:5000, Jackson ImmunoResearch).

### Quantitative RT-PCR

Total RNA was collected using Trizol (Invitrogen) followed by cleanup using RNA Clean and Concentrator (Zymo) using the manufacturer’s protocol. cDNA was obtained with reverse transcription using SuperScript III kit (Invitrogen). qPCR was performed using primers list below (SYBR, Invitrogen)

A list of primers used in the paper

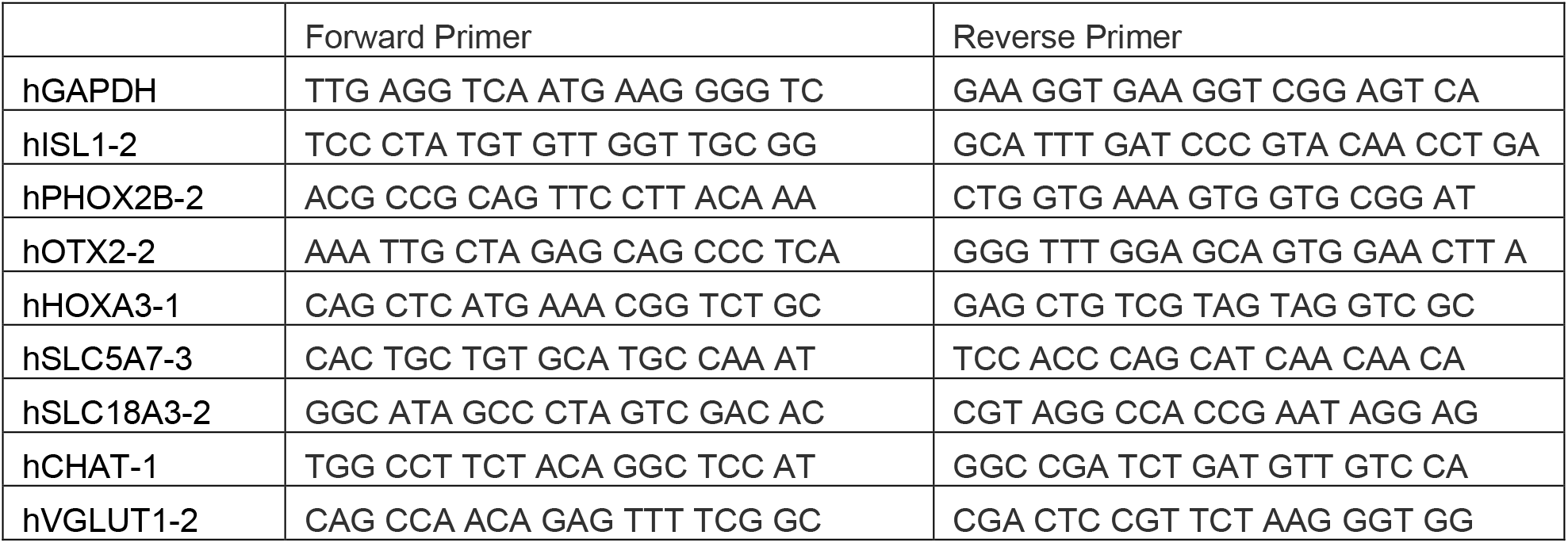

### Immunofluorescence Staining

The cells were fixed by adding PFA for 15 minutes. The cells were then washed in 1.5 mL of PBS and then lysed and blocked in a blocking buffer containing PBS, 0.1%Triton-X, 0.1% NaN3, and 5% CCS for 30 minutes. The cells were stained with the corresponding primary antibodies diluted in blocking buffer 30 minutes at room temperature or 4C overnight. The cells were then washed three times with blocking buffer before addition of the corresponding secondary antibody in blocking buffer for 1 hour. The cells were then washed three times with blocking buffer before the addition of DAPI (Invitrogen) in PBS for 1 minute. The cells were washed three times and then imaged. Antibodies used: mouse-ISL1 (1:100, DSHB), goat PHOX2B (1:100, Santa Cruz), rabbit-TUJI (1:1000, Covance), goat OTX2 (1:100, R&D), mouse HOXA3 (1:500, R&D), mouse TUJI (1:1000, Covance), mouse MAP2 (1:500, Sigma), rabbit VGLUT1 (1:1000, Synaptic system) and mouse Flag (1:1000, Sigma).

### Image analysis

For the analysis of the IF staining MetaMorph^®^ Microscopy Automation & Image Analysis Software was used. To properly analyze the cells a minimum width of 15μm and a maximum of 30μm was used. The DAPI channel was used as a positive control and the intensity above the local background was manually adjusted for each sample such that only true positives were considered. After computation the nuclear intensity and counts were checked manually for any false positives or counts which included more than one cell per count.

### Electrophysiology

Recordings of the intrinsic and active membrane properties were performed as previously described (Zhang et al., 2013). Briefly, cells were recorded in a bath solution containing (in mM): NaCl 140, 10 HEPES, 10 Glucose, 5 KCl, 3 CaCl2, 1 MgCl_2_ (pH adjusted to 7.4, 300 Osm)., along with 1 μM Tetrodotoxin (American radiolabeled chemicals). 10 μM CNQX (Tocris) was applied to block α-Amino-3-hydroxy-5-methyl-4-isoxazolepropionic acid (AMPA)-receptor excitatory postsynaptic currents (EPSCs), The miniEPSC recordings were performed in voltage-clamp mode with internal solution containing (in mM): 135 CsMeSO3, 10 HEPES, 8 NaCl, 0.25 EGTA, 2 MgCl_2_, 4 MgATP, 0.3 Na2GTP, 5 PO4-creatine, 1 QX134 (pH adjusted to 7.3, 304 Osm).

## Datasets used

scRNA sequencing for H9 cells used: SRR2977655, SRR2977656, SRR2977657, SRR2977658, SRR2977659, SRR2977660, SRR2977661, SRR2977662, SRR2977663, SRR2977664, SRR2977665.

---

**Fig. S1 – supplementary to Figure 1.**
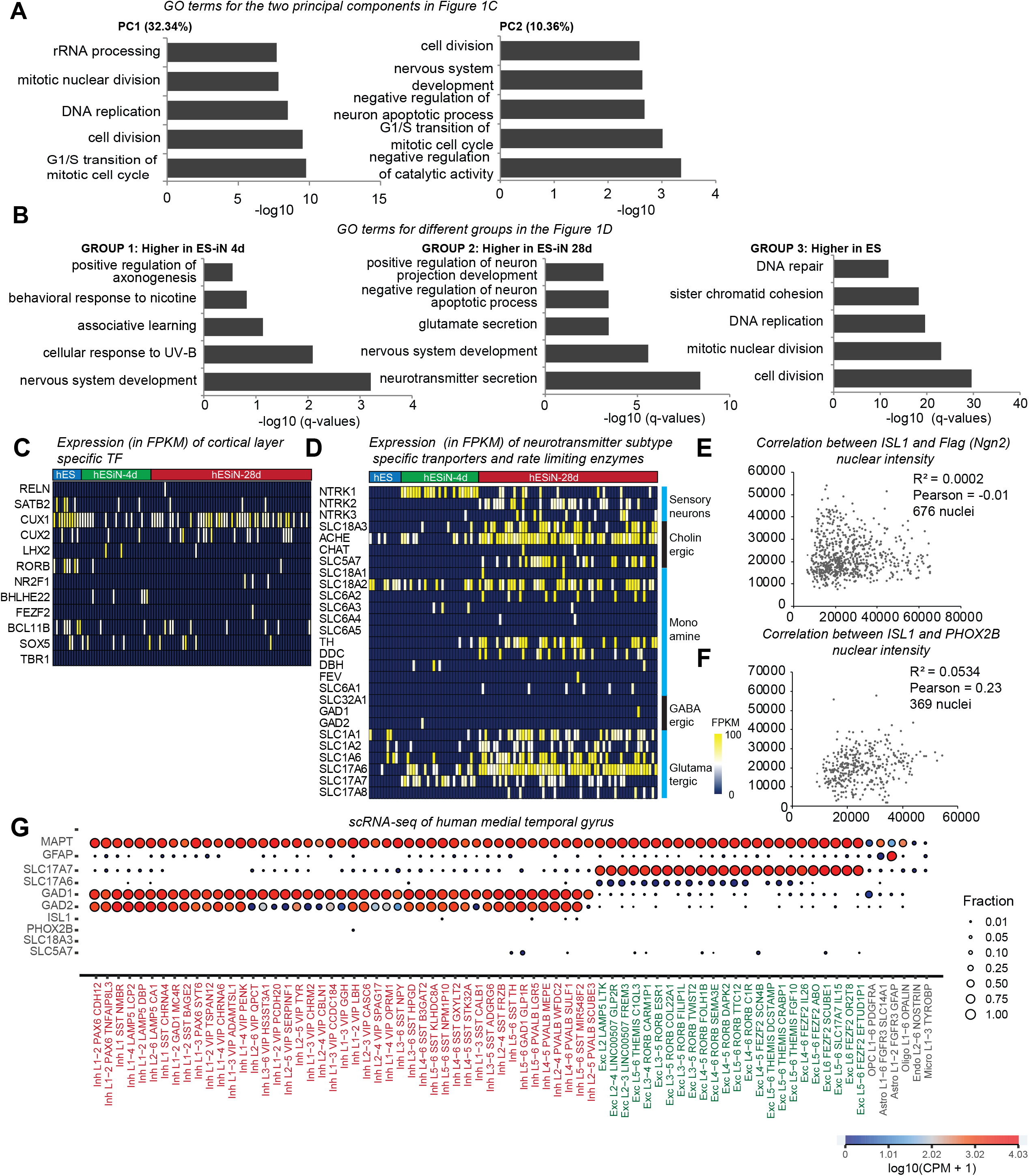
(A) The gene ontology terms for genes enriched in the first and second principal component shown in Figure 1C. (B) The gene ontology terms for genes enriched in the three groups described in Figure 1D (C) Heatmap showing the expression of different cortical layer markers during the process of reprogramming. (D) Heatmap showing the expression different transporters and channel proteins for different neurotransmitter subtype markers during the process of reprogramming. (E) Correlation plot of the nuclear intensity of ISL1 and FLAG (NGN2) of 676 nuclei of day 4 NGN2 iN cells. There is no observed correlation between the two variables. (F) Correlation plot of the nuclear intensity of ISL1 and PHOX2B of 369 nuclei of day 4 NGN2 iN cells. There is no observed correlation between the two variables. (G) Single cell RNA-seq data from primary human neurons reveals little co-expression of excitatory markers with cholinergic genes in cortex. Graph showing the fraction of cells and its corresponding log(CPM+1) value for the individual population of neurons and non-neuronal cells in human medial temporal gyrus. Data was obtained from celltypes.brain-map.org.

**Fig. S2 – supplementary to Figure 2.**
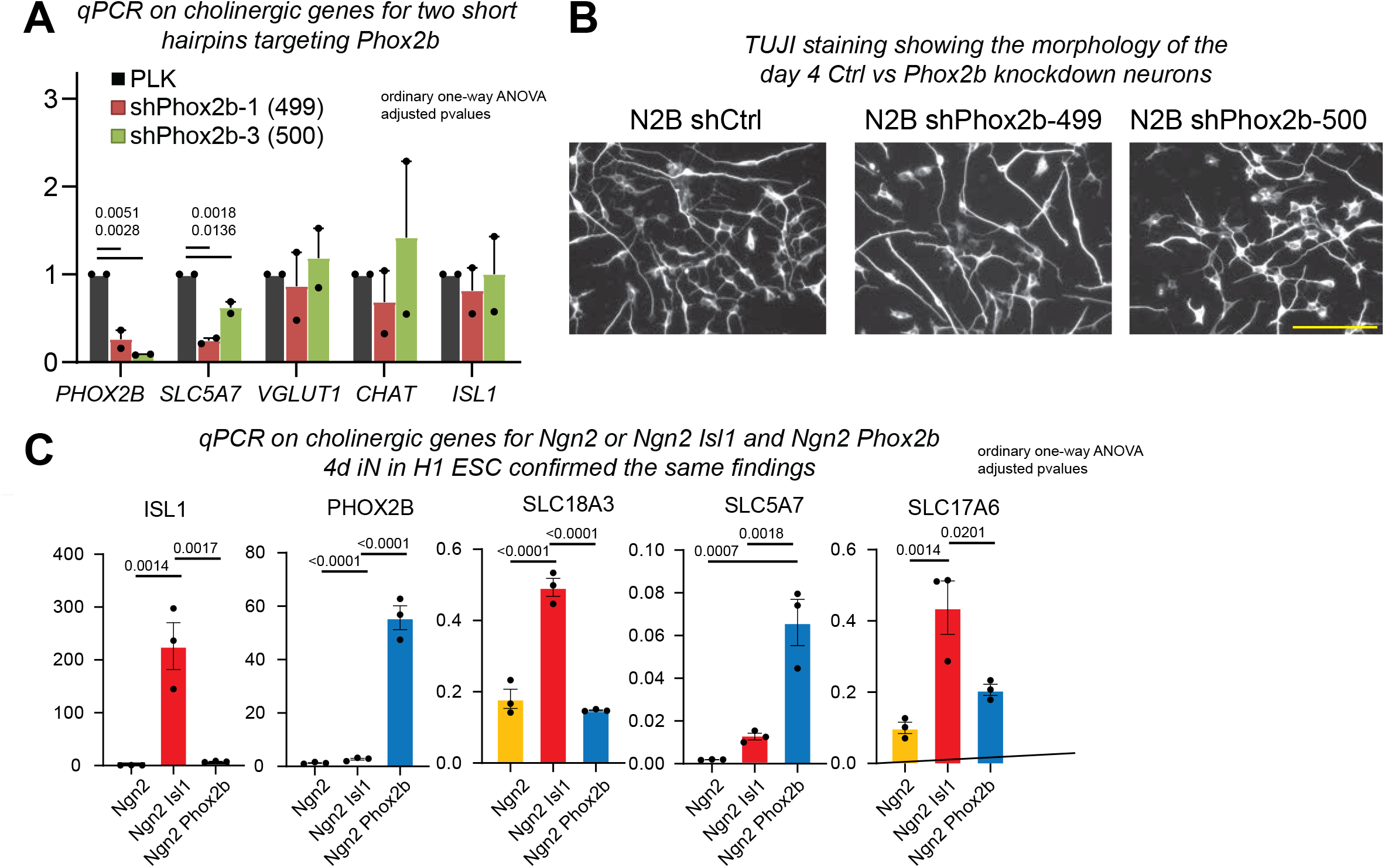
(A) Quantitative RT-PCR examining the day 4 NGN2 iN cells infected with a control, or two different Phox2b short hairpins (termed 499 and 500). Shown is expression for PHOX2B, two cholinergic markers (CHAT, SLC5A7), transcription factor (PHOX2B, ISL1) and a glutamatergic marker (vGLUT1). 499 and 500 denotes hairpin id outlined in the method section (N=3, error bars = s.e.m. ANOVA. Exact adjusted p-values are shown in the graph) (B) Immunofluorescence images of β-III-tubulin staining of day 4 NGN2 iN cells infected with a control, Phox2b-499 or Phox2b-500 short hairpin virus. (C) Quantitative RT-PCR examining the day 4 NGN2 iN cells alone or coexpressing ISL1 and PHOX2B derived from another human embryonic stem cell line (H1 hESC) for three cholinergic markers (SLC18A3, SLC5A7) and a glutamatergic marker (vGLUT1). (N=3, error bars = s.e.m. Statistical testing was performed using ANOVA. Exact adjusted p-values are indicated in the graph)

**Fig. S3 – supplementary to Figure 3.**
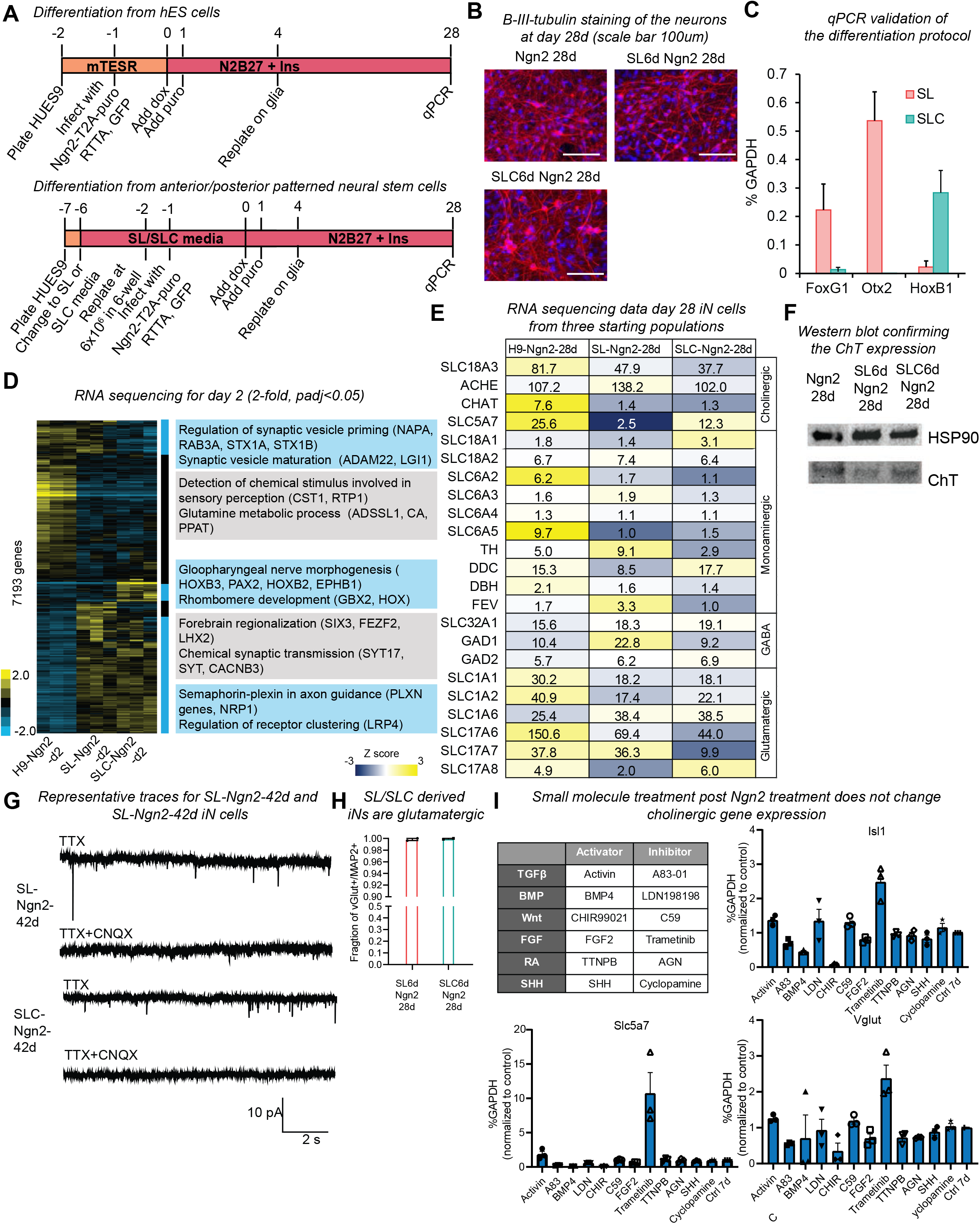
(A) Schematics showing (top) the conventional protocol to differentiate iN cells from hES cells and differentiation of iN cells from anterior and posterior neural stem cells (bottom). (B) Immunofluorescence images of β-III-tubulin staining of day 28 NGN2 iN cells generated with the conventional protocol from hES cells and from anterior and posterior neural stem cells. Scale bar=100μm. (red: β-III-tubulin and blue: DAPI) (C) Quantitative RT-PCR validation of the expression of anterior (*FOXG1, OTX2*) and posterior (*HOXB1*) genes in neural progenitor cells patterned with SL or SLC. Error bars = s.e.m. (D) Hierarchical clustering of differentially expressed genes (>2-fold change and p adj<0.05) two days after NGN2 expression in the three starting populations (H9, SL and SLC). Significant gene ontology terms and the genes contributing to them for each highlighted cluster are listed (p adj<0.05). (E) A summary of the regional identity and subtype specific marker expression for each of the three populations (H9-NGN2-28d, SL-NGN2-28d, SLC-NGN2-28d). The box was colored using the z-normalized log2(TPM+1) and the number of the box represents the TPM+1 value. Note that SL-NGN2-28d, SLC-NGN2-28d silenced majority all other neurotransmitter programs albeit incomplete. (F) Western blot analysis of whole cell lysates from day 28 iN cells derived from H9, SL or SLC (pre-differentiated and patterned for 6 days) using the ChT and HSP90 antibodies. (G) Representative traces of miniature excitatory postsynaptic currents (EPSCs) for SL-NGN2-42d and SLC-NGN2-42d iN cells in the presence of the Na^+^ channel blocker tetrodotoxin (TTX) (top traces) or both TTX and the AMPA receptor blocker CNQX (bottom traces). (H) Graph showing the fraction of VGLUT1 among MAP2^+^ iN cells at day 28. In both conditions > 90% of MAP2^+^ neurons expressed the excitatory marker vGluT. Error bars = s.e.m. (N=2, 10 visual fields per replicate) (I) qRT-PCR gene expression analysis for Isl1, Scl5a7, and vGluT of Ngn2 iN cells treated for the first 7 days in the presence of small molecules indicated. Small molecules represent activators and inhibitors of the signaling pathways shown in the table (top left). CHIR99021 produced only immature neurons and Trametinib induced significant cell death. Error bars = s.e.m.

**Fig. S4 – supplementary to Figure 4.**
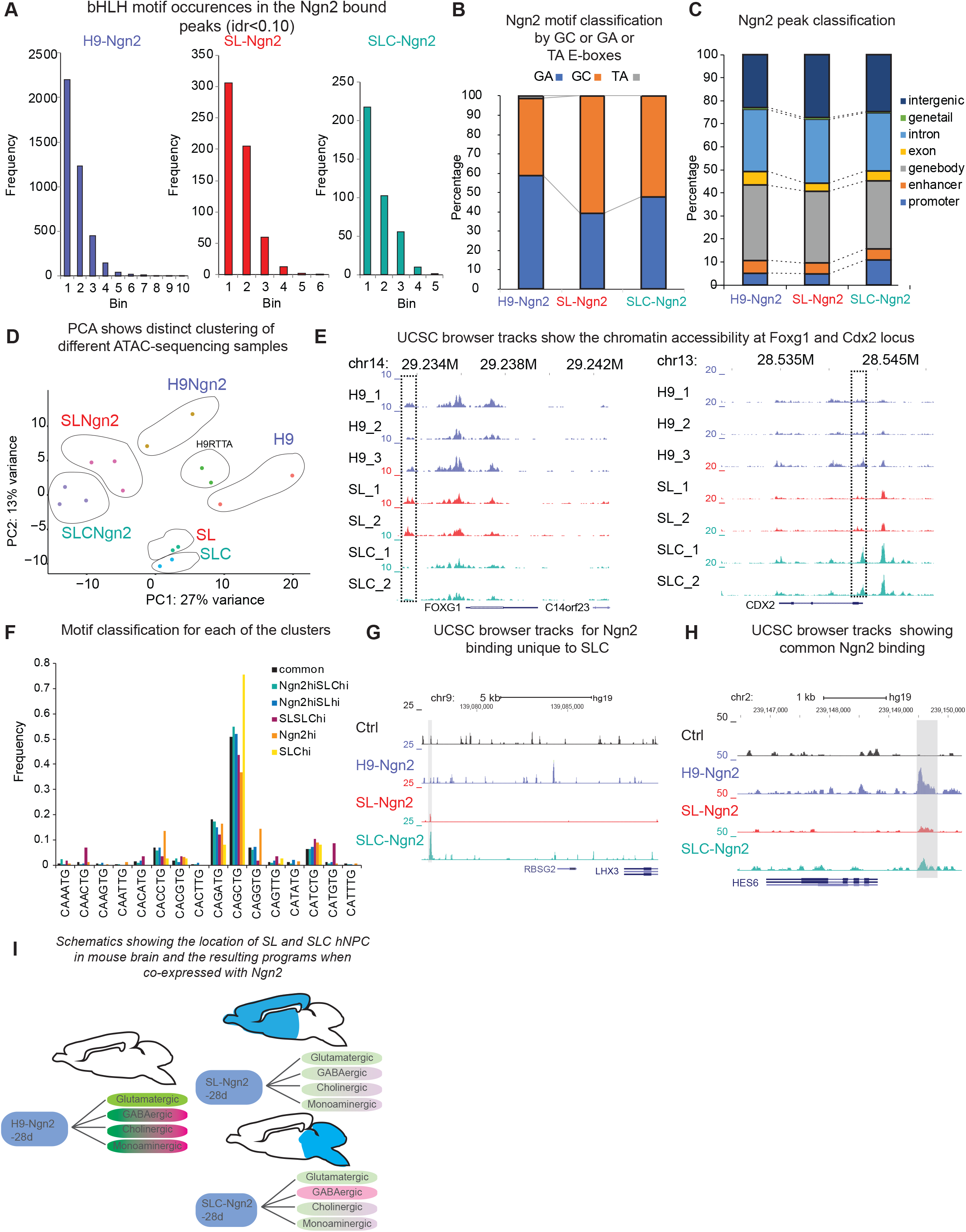
(A) bHLH motif occurrence in each consensus NGN2 ChIP-sequencing peak (idr q<0.10, n=2) for H9-NGN2, SL-NGN2 and SLC-NGN2. The NGN2 ChIP peaks with one bHLH motif were most frequent followed by 2,3,4, and 5 motifs in an exponentially decreasing manner. (B) Types of E-box motif classification (GC, GA or TA) enriched under NGN2 ChIP-seq peaks in ES cells (H9), anterior (SL), and posterior (SLC) neural progenitors 2 days after NGN2 infection. The vast majority of E-box motifs found are either of the GA or GC type in all three conditions. (C) Genomic distribution of NGN2 peaks is similar between ES cells (H9), anterior (SL), and posterior (SLC) neural progenitors 2 days after NGN2 infection. Promoter = 2000bp upstream and 1000bp downstream of TSSs, enhancer = 2000-10000bp upstream of TSSs, the remaining sequences are classified as intergenic. (D) Principal component analysis of ATAC sequencing for the three starting cell populations (H9, SL and SLC) with and without NGN2 expression for 2 days. Note that all six populations cluster and their replicates cluster closely in groups. (E) Genomic browser tracks showing ATAC sequencing signals for the three starting populations (H9, SL and SLC) in the *FOXG1* (left) and *CDX2* (right) locus. The dotted boxes highlight the differentially accessible areas. (F) E-box motif classification for each of the clusters in Figure 4A. No obvious difference in E box motifs among the different groups (G) Genomic browser tracks showing a NGN2 binding site highly enriched in SLC neural progenitor cells (slim grey highlight) far upstream of the *RBSG2/LHX3* locus. (H) Genomic browser tracks showing a shared NGN2 site in H9-hES cells, SL and SLC neural progenitor cells at the *HES6* locus. (I) Schematic showing the regional identity of the SL (anterior) and SLC (posterior) neural progenitor cells (highlighted in blue) projected onto the mouse brain. The tree highlights the neurotransmitter subtype programs in the resulting d28 iN cells from SL (anterior) and SLC (posterior) neural progenitor cells. Green represents full, green-red partial, and red represents no neurotransmitter programs. The color intensity represents the level of expression based on the table in Figure S3E. Compared to H9-NGN2 28d iN cells, iN cells from SL and SLC have global downregulation of GABAergic, monoaminergic and cholinergic genes (Figure S3E). The resulting SL and SLC d28 iN cells still have partial neurotransmitter programs, however.

**Fig. S5 – supplementary to Figure 5.**
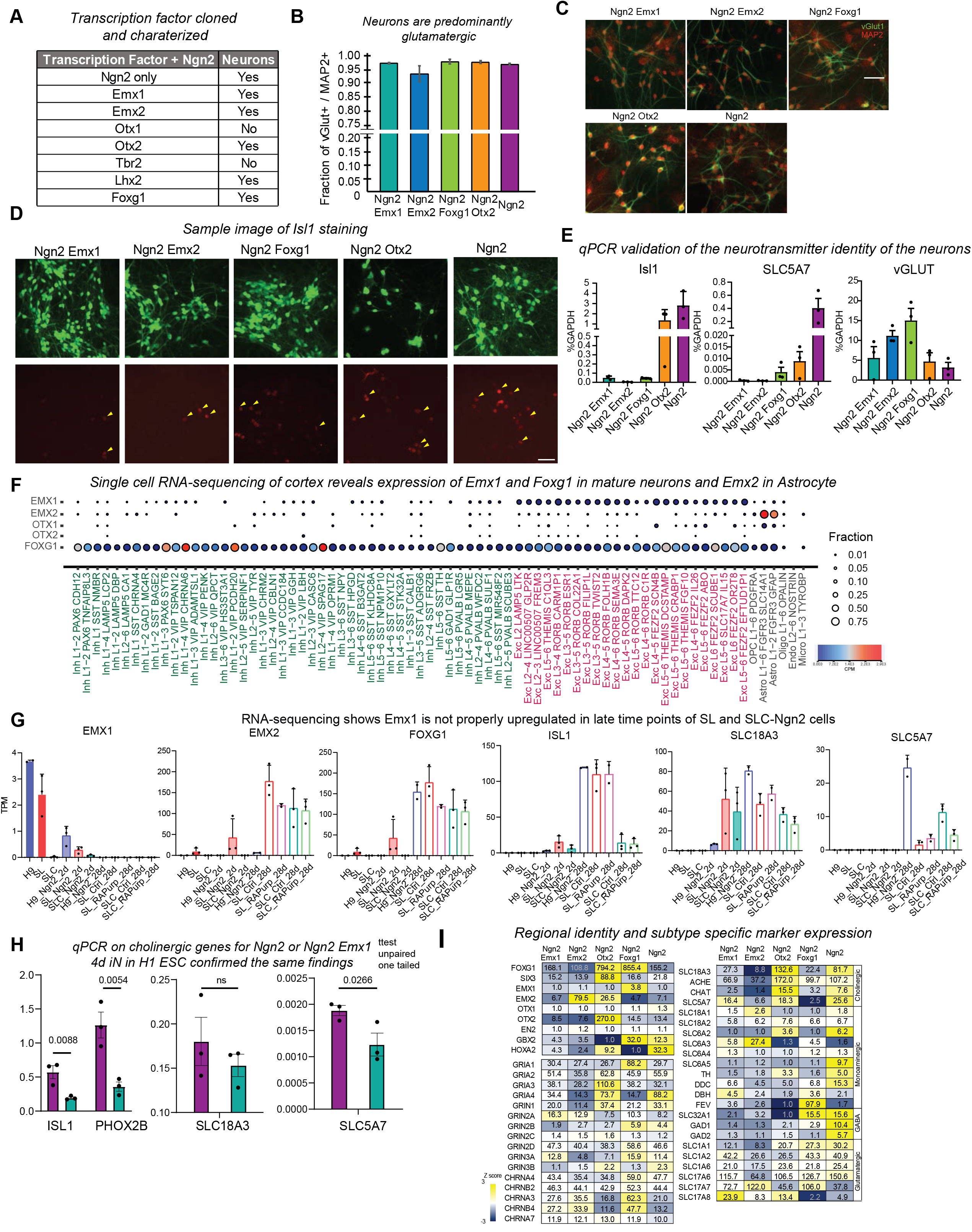
(A) Table summarizing the results of the transcription factors co-expressed with NGN2. With the exception of OTX1 and TBR2, all transcription factors produced neurons with NGN2 at day 28. (B) Graph showing the fraction of VGLUT1 among MAP2 positive iN cells at day 28 generated with NGN2 alone or in combination with EMX1, EMX2, FOXG1, and OTX2. All conditions generated > 90% vGluT^+^ neurons. Error bars = s.e.m. (C) Representative immunofluorescence micrographs of neurons generated with indicated transcription factors for vGluT1 (green) and MAP2 (red). Scale bar: 50μm (D) Representative images of GFP (top) and Isl1 (bottom) immunofluorescence for iN cells generated with indicated transcription factors. Scale bar: 50μm (E) Quantitative RT-PCR validation of the expression of *ISL1, SLC5A7* and *VGLUT1* in iN cells generated with indicated transcription factor combinations. Error bars = s.e.m. (F) Single cell RNA expression of *EMX1, EMX2, OTX1, OTX2 and FOXG1* in cells from human medial temporal gyrus. The circle size indicates the fraction of cells expressing the gene and the color indicates expression level of indicated group of cells measured by logCPM+1 value. Data was obtained from celltypes.brain-map.org. (G) RNA sequencing data for indicated genes in H9, SL and SLC before and 2 days and 28 days after NGN2 expression. The day 28 marked with RAPurp were treated for retinoid acid and purmo-8 erphamine after plating on glia to day 28. Error bars = s.e.m. (H) Quantitative RT-PCR examining the day 4 NGN2 iN cells alone or co-expressing EMX1 derived from another cell line (H1 hESC) for two cholinergic markers (SLC18A3, SLC5A7), ISL1, PHOX2B and a glutamatergic marker (vGLUT1). N=3; Student’s t-test, unpaired, one tailed; p values shown above histograms; error bars = s.e.m. (I) A summary of the regional identity and subtype specific marker expression for each of the five different kind of iN cells generated with the transcription factor combination indicated. The boxes are colored using the z-normalized log2(TPM+1) and the number in the boxes shows the TPM+1 value. Note the efficient silencing of cholinergic, monoaminergic, and GABAergic genes in NGN2:EMX1 iN cells.

**Fig. S6 – supplementary to Figure 6.**
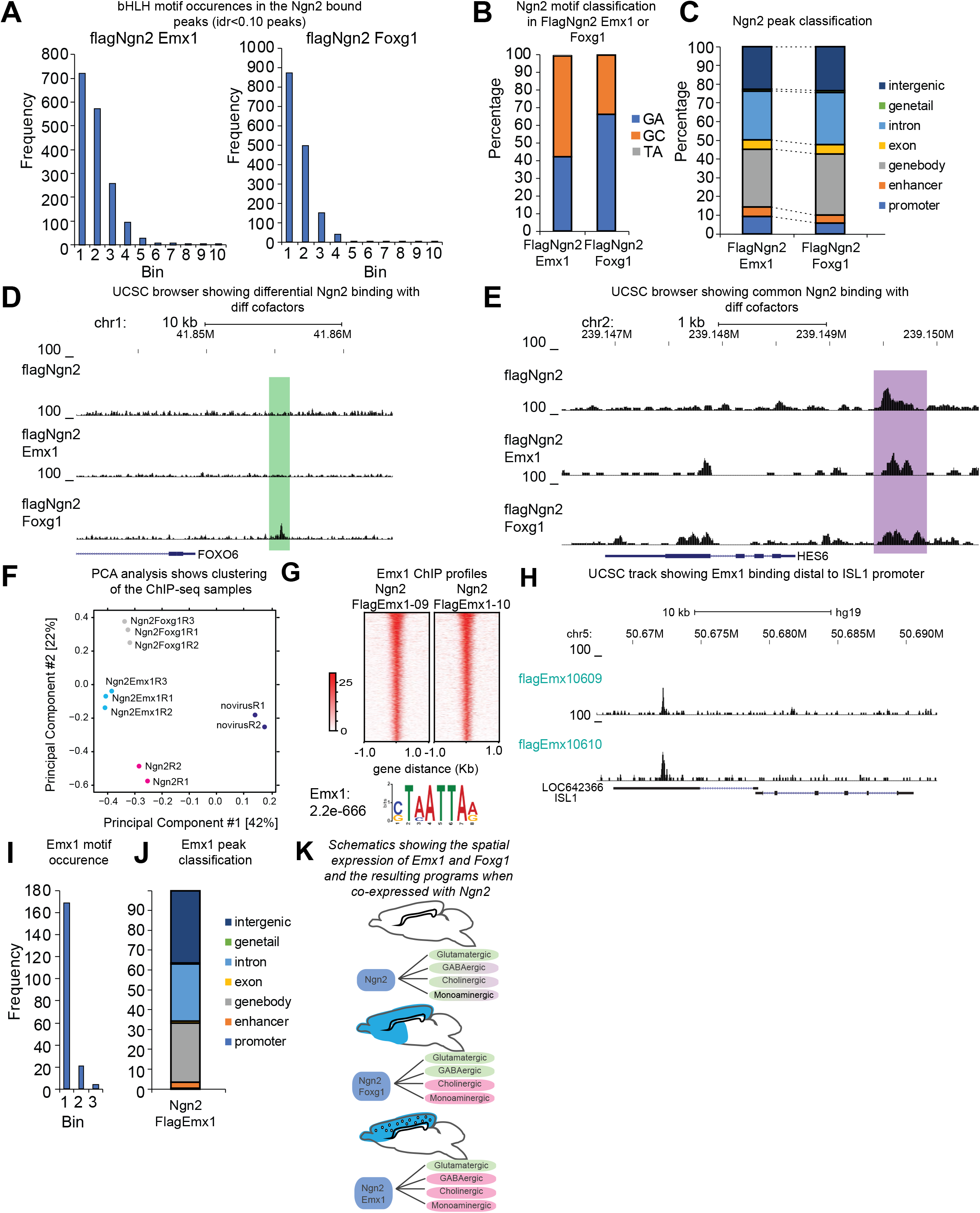
(A) Distribution of NGN2 ChIP-seq peaks based on the number of bHLH motifs under each peak (idr q<0.10, n=2) in cell infected with flagNGN2:EMX1 or flagNGN2:FOXG1. Peaks with one bHLH motif were most abundant in both cases and the frequency distribution decreases exponentially with the number of motifs within each peak. In flagNGN2:EMX1 cells there were more peaks that contained more than one motif whereas in flagNGN2:FOXG1 there were more peaks that contained only one motif than multiple. (B) Types of E-box motif classification (GC, GA or TA type) for NGN2 peaks in cells infected with NGN2:EMX1 or NGN2:FOXG1. The tow most prominent E-box motif types by far are the GA and GC types with a preference of the GC type in flagNGN2:EMX1 cells and a preference of the GA type in flagNGN2:FOXG1 cells. (C) Genomic peak distribution of NGN2 peaks in ES cells infected with either NGN2:EMX1 or NGN2:FOXG1. Promoter = 2000bp upstream and 1000bp downstream of TSSs, enhancer = 2000-10000bp upstream of TSSs, the remaining sequences are classified as intergenic. (D) Genomic browser tracks of the human *FOXO6* locus showing the flagNGN2 ChIP-seq signal in ES cells infected with flagNGN2, flagNGN2:EMX1, or flagNGN2:FOXG1. The light green box highlights the NGN2 peak unique to the cells expressing flagNGN2:FOXG1. (E) Genomic browser tracks of the human *HES6* locus showing the ChIP-seq signal in the three cell populations infected with flagNGN2, flagNGN2:EMX1 or flagNGN2:FOXG1. The purple box highlights the NGN2 peak common in all three populations. (F) Principal component analysis of the ChIP samples (novirus Ctrl, NGN2, NGN2:EMX1 and NGN2:FOXG1). R1-3=replicate samples. (G) *Top:* Heatmaps of two replicates of the flagEMX1 ChIP-seq profile in NGN2:flagEMX1 infected ES cells. ChIP-seq peaks are displayed +/− 1kb from the peak summit. *Bottom:* The EMX1 motif was significantly enriched among the sequences of the significantly called EMX1 ChIP-seq peaks. (H) Genomic browser tracks showing an EMX1 peak upstream of the *ISL1* promoter in both replicates (0609 and 0610 denote replicates). (I) Distribution of EMX1 ChIP-seq peaks based on the number of EMX1 motifs under each peak (idr q<0.10, n=2) in ES cells infected with NGN2:flagEMX1. Unlike for NGN2 peaks and E-box motifs (Figure S6A), EMX1 peaks had predominantly a single EMX1 motif. (J) Genomic classification of EMX1 peaks in ES cells infected with NGN2:EMX1. Promoter = 2000bp upstream and 1000bp downstream of TSSs, enhancer = 2000-10000bp upstream of TSSs, the remaining sequences are classified as intergenic. (K) Schematic outlining the expression pattern of EMX1 and FOXG1 (highlighted in blue) projected onto the mouse brain. Note that EMX1 is not expressed in inhibitory neurons (denoted by blank circles). The trees below the drawings indicate the neurotransmitter subtype programs in the resulting d28 iN cells from NGN2, NGN2:EMX1 and NGN2:FOXG1 expression in human ES cells (green represents full, green-red partial, and red represents no neurotransmitter programs).

